# Accounting for longitudinal peak quality metrics with MSstats+ enhances differential analysis in proteomic experiments with data-independent acquisition

**DOI:** 10.1101/2025.09.11.675573

**Authors:** Devon Kohler, Eralp Dogu, Mrittika Bhattacharya, Ozge Karayel, Manuel Magana, Anthony Wu, Veronica G Anania, Olga Vitek

## Abstract

Mass spectrometry-based proteomics with data-independent acquisition benefits from advanced instrumentation and computational analysis. Despite continued improvements, the quality of quantification may be poor for some measurements. As the scale of proteomic experiments increases, these poor-quality measurements are challenging to characterize by hand, yet they undermine the detection of differentially abundant proteins and the downstream biological conclusions. We introduce *MSstats+*, a computational workflow that takes as input not only peak intensities reported by tools such as Spectronaut, but also quality metrics such as peak shape and retention time, and longitudinal run order profiles of these metrics. *MSstats+* translates these metrics into a single measure of quality, and downweights poor quality measurements when detecting differentially abundant proteins. The method offers a natural treatment of missing values, weighting the imputed values according to the quality metrics in the run. We demonstrate the accuracy of the resulting differential analysis in four experiments: two custom benchmarking studies with intentionally induced anomalies, a controlled mixture of proteomes, and a large-scale clinical investigation. *MSstats+* is implemented in the family of open-source R/Bioconductor packages *MSstats*.

## 1. Introduction

Advances in data-independent acquisition (DIA) mass spectrometry (MS)-based proteomics hold high promises for both basic and clinical research [1–4]. Increasingly larger in scope, modern investigations simultaneously quantify thousands of proteins in thousands of samples [5, 6]. Novel instrumentation has improved acquisition speed, sensitivity, and ion manipulation, leading to deeper and more consistent measurements [7, 8]. DIA strategies have reduced stochasticity and improved detection of low-abundant ions [9–11]. Spectral processing tools such as Spectronaut [12], DIA-NN [13], Skyline [14, 15], and FragPipe [16], identify and quantify peptide ions and their fragments with stringent false discovery rate (FDR) control to retain high-confidence identifications [17–20]. The tools report intensities of the identified peaks as the area under the curve or the height of the peak at apex.

Despite great advances, low-quality quantitative measurements and missing values remain. Low-quality measurements can be caused by a variety of factors, such as peak interference, poor chromatographic shape, or misidentification [21]. Peak intensities may be entirely missing when the abundance of the analyte is below the limit of detection of the instrument, a situation often referred to as missing not at random (MNAR). Missing peak intensities may also be introduced stochastically, for reasons such as peak picking errors or fragment interference [22], and are referred to as missing at random (MAR) or missing completely at random (MCAR) [23, 24]. The extent of low-quality and missing measurements is exacerbated in large-scale experiments that profile many samples and many analytes [5, 25]. Advanced statistical methods that take as input the intensities of the identified spectral peaks, summarize these intensities into protein-level abundances in a run, and model the stochastic structure of the abundances to detect differentially abundant proteins, must take the varying quality of measurements into account to support effective use.

Numerous statistical methods and tools have been developed for protein-level summariza-tion and differential analysis. The methods differ both in scope and in the underlying models and assumptions. Some focus solely on protein-level summarization (e.g., *MaxLFQ* [26]), others focus only on statistical modeling of protein-level abundances (e.g., *limma* [27], *DEqMS* [28], *ROTS* [29]), and still others offer end-to-end workflows (e.g., *MSstats* [30], *limpa* [31], *msqrob2* [32, 33]). Some methods have advanced modeling features, such as characterization of uncertainty in protein summarization in *limpa* and Empirical Bayes in *DEqMS, limma* and *limpa* that can positively impact their performance. In all cases, the most common strategy to distinguish high- and low-quality measurements in differential analysis is feature selection [34]. In the context of DIA, a feature refers to a fragment ion.

Feature selection, such as the selection of a subset of fragment ions with the highest average abundance, or its more advanced implementation [35–37], relies on the fact that poorer quality measurements are associated with lower intensities, larger number of missing values, and/or higher variation. Although effective in some cases, this strategy introduces the risk of circular analysis (also called “data leakage” or “double dipping”)[38, 39], where intensities are used to select features and then those same intensities are used in differential analysis. For example, **Supplementary Fig. 1** uses a controlled mixture from Navarro et al. [40] to show that selecting highest intensity features and subsequently testing those same features for differential abundance inflates confidence in the results and increases false discoveries. Moreover, missing values may survive feature selection. Determining their causes is nontrivial, and their statistical treatment such as imputation is an active area of research [22, 41, 42].

**Figure 1.**
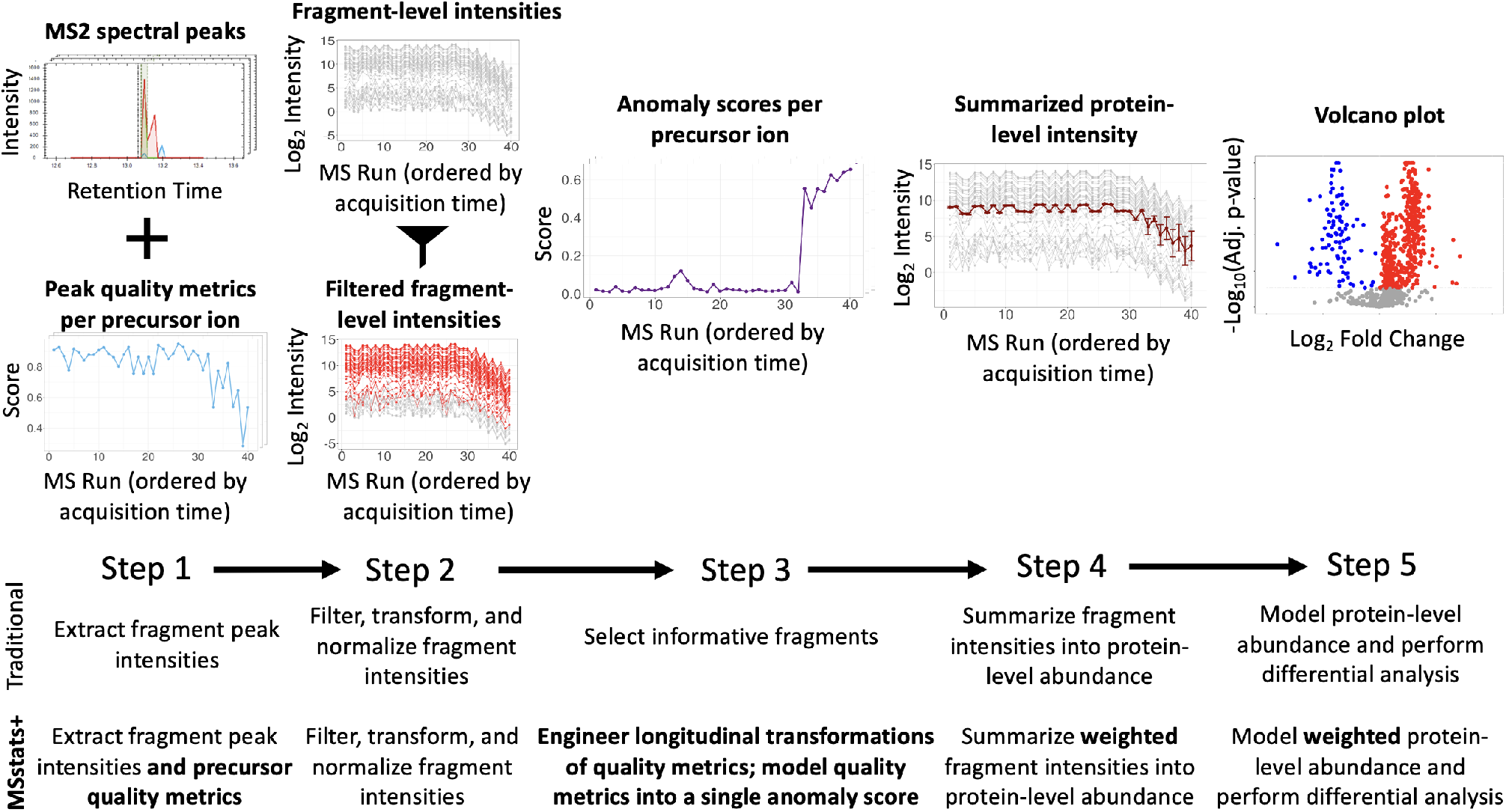
*MSstats+* integrates spectral peak quality metrics into differential analysis. *MSstats+* is illustrated using precursor LATQSNEITIPVTFESR.3 and protein P04792 in the K562 benchmark, and contrasted with the traditional workflow.

The difficulties with low-quality and missing measurements stem from the fact that, feature intensities are traditionally the only input made available to differential analysis. Yet data processing tools report metrics such as delta retention time, peak width, mass accuracy, and other scores that are complementary to peak intensities and are informative of the quality of identification and quantification. Until recently, the use of these metrics was confined to longitudinal profiling for system suitability and quality control, to characterize analytes in controlled mixtures interleaved between the samples of interest, or spiked directly into the samples as standards [21, 43–45]. However, these metrics are typically not considered as part of the differential analysis of endogenous proteins of interest.

Recent studies have highlighted the value of such metrics for peptide and protein quantification. QuantUMS [46] uses the similarity of elution profiles between a peptide and a “best” fragment, and downweights measurements with inconsistent profiles when performing protein-level summarization. However, the implementation is specific to DIA-NN, and requires prior knowledge of the “best” fragment. Similarly, MCQR [47] uses the distributions of retention time alignment and chromatographic peak widths to filter out poor quality measurements; however, it requires manual tuning of user-defined cutoffs, which can be inconsistent between precursors and is not scalable to large and complex datasets (**Supplementary Fig. 2**).

**Figure 2.**
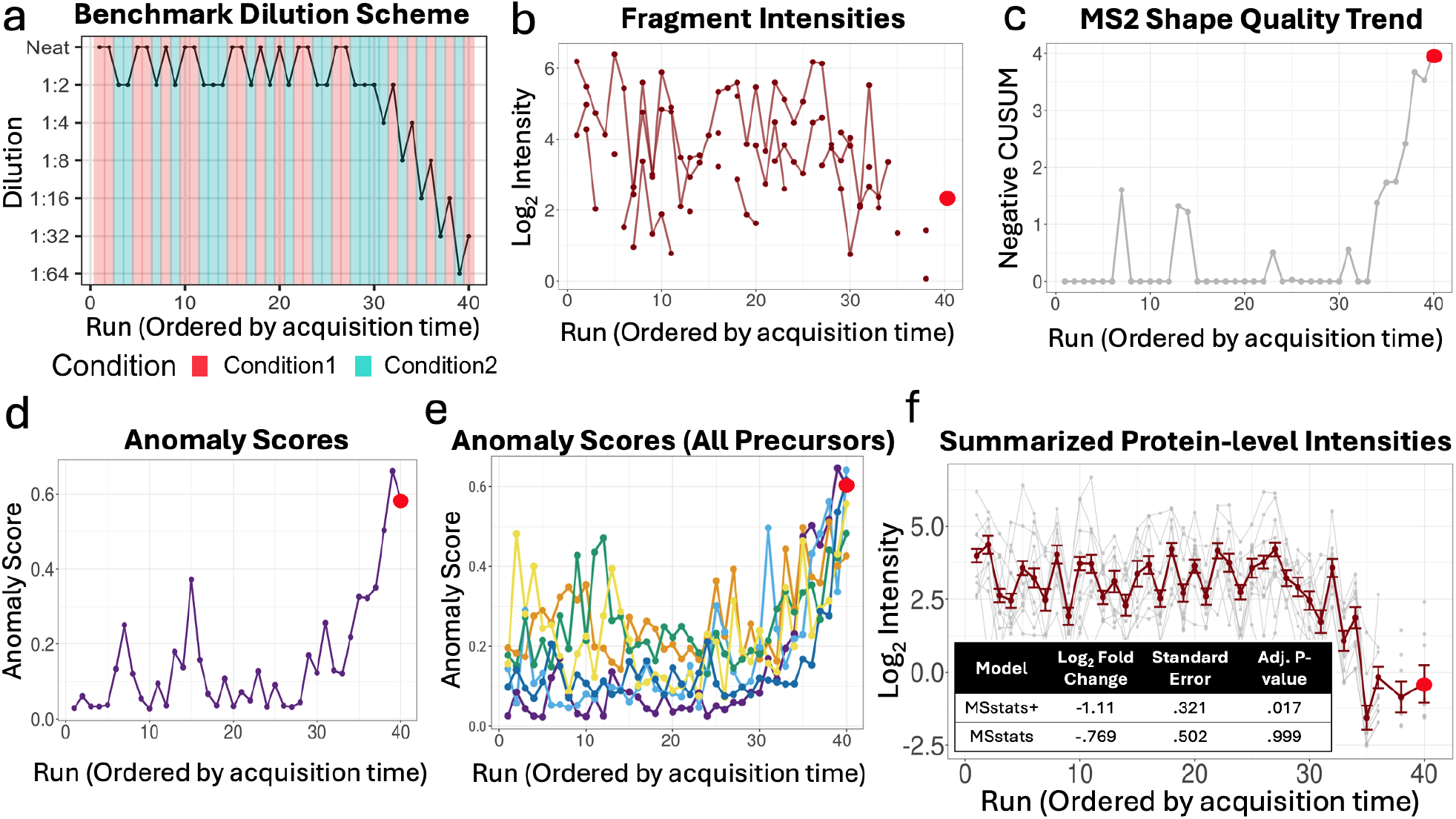
K562 benchmark, protein A4D126: *MSstats+* corrected for poor quality measurements in differential analysis. Run 40 is indicated with a red dot. (**a**) Dilution scheme for the K562 and CSF benchmark experiments. Different dilution amounts were used to mimic two experimental conditions with a log_2_ fold change of 1. Lower amounts of material were injected over the last 10 runs to mimic instrumental drift (while preserving the fold change). (**b**) The precursor had 3 fragments, and only one fragment measured in run 40. The log_2_ intensity for the measured fragment in run 40 was unremarkable. (**c**) Precursor ALAEDQINSK.2 showed a decrease in MS2 shape quality score in later acquired runs. Negative CUSUM had individually high jumps in runs 7, 13, and 14, and a sustained increase across runs 34-40. (**d**) This deviation contributed to the high overall anomaly score for that precursor in run 40. (**e**) Anomaly scores across all the precursors of A4D126 were higher in the later runs, similarly to run 40, reflecting the intentionally introduced anomalies. Colors indicate different precursors of the protein. (**f**) Protein-level summarization of A4D126 with *MSstats+* accounted for the uncertainty in the quantification of the later runs such as run 40. The dark red line is the protein-level summary. The gray lines are the fragments. Error bars are standard deviations of the run-level estimates. *MSstats+* produced a log_2_ fold change closer to the ground truth (true value: –1), and reduced the standard error by 36%.

This manuscript introduces *MSstats+*, a comprehensive statistical workflow for integrating quality metrics of endogenous peaks into the differential analysis. *MSstats+* considers not only the values of the quality metrics, but also their longitudinal profiles such as isolated deviations in quality due to interference, or systematic deviations in quality due to instrumental drift that are monitored in system suitability and quality control but are overlooked by feature selection. *MSstats+* uses unsupervised isolation forest [48] to summarize this information into a single anomaly score per peptide peak and downweights anomalous intensities in the differential analysis workflow. By weighting the peaks according to information that is complementary to peak intensities, as opposed to peak intensities directly, *MSstats+* separates quality assessment from statistical modeling and avoids double dipping. The approach naturally incorporates missing values after imputation by weighting the imputed values according to their anomaly scores.

We evaluated *MSstats+* on three controlled experiments with known ground truth: two ad-hoc studies with intentionally introduced anomalies and a previously published mixture of proteomes [49]. We compared the performance of *MSstats+* to that of *limma* [27], *limpa* [31], *msqrob2* [32, 33], *DEqMS* [28], and *MSstats* [30, 50] in terms of sensitivity and specificity to detect differentially abundant proteins, and demonstrated the advantage of including peak quality considerations in the analysis. To demonstrate the real-world relevance and scalability of *MSstats+*, we applied it to a clinical cohort [51] and demonstrated better control of poor quantitative measurements.

*MSstats+* is implemented as part of the family of open-source R/Bioconductor packages *MSstats* [52–54] making it accessible for routine and modular use. The source code and documentation are hosted at github.com/Vitek-Lab/MSstats, with planned released on Bioconductor (October 2025).

## 2. Results

### 2.1. Overview of *MSstats+*

Traditional differential analyses of DIA-MS proteomic experiments take as the sole input intensities of identified and quantified fragment ion peaks. However, upstream data processing tools, such as Spectronaut, provide additional metrics (e.g., delta retention time, mass accuracy, peak width, peak shape, etc.) that capture peak quality in ways that are complementary to the intensities. For example, non-Gaussian peak shape or differences in predicted versus observed retention time can indicate unreliable quantification even when intensity values are high. Conversely, peaks with low intensities may be informative if the quality metrics are stable.

*MSstats+* leverages quality metrics in the differential analysis workflow as illustrated in **Figure 1** and detailed in **Section 4.2**. In Step 1, the peak intensities and peak quality metrics are extracted from the spectral processing tool. **Supplementary Fig. 3** illustrates this in the case of Spectronaut. Spectronaut reports peak quality metrics at multiple levels, specifically elution group, fragment group (precursor ion), and individual fragments. While *MSstats+* uses fragment peak intensities to maximize accuracy of quantification, it uses precursor quality metrics for reasons of computational efficiency.

**Figure 3.**
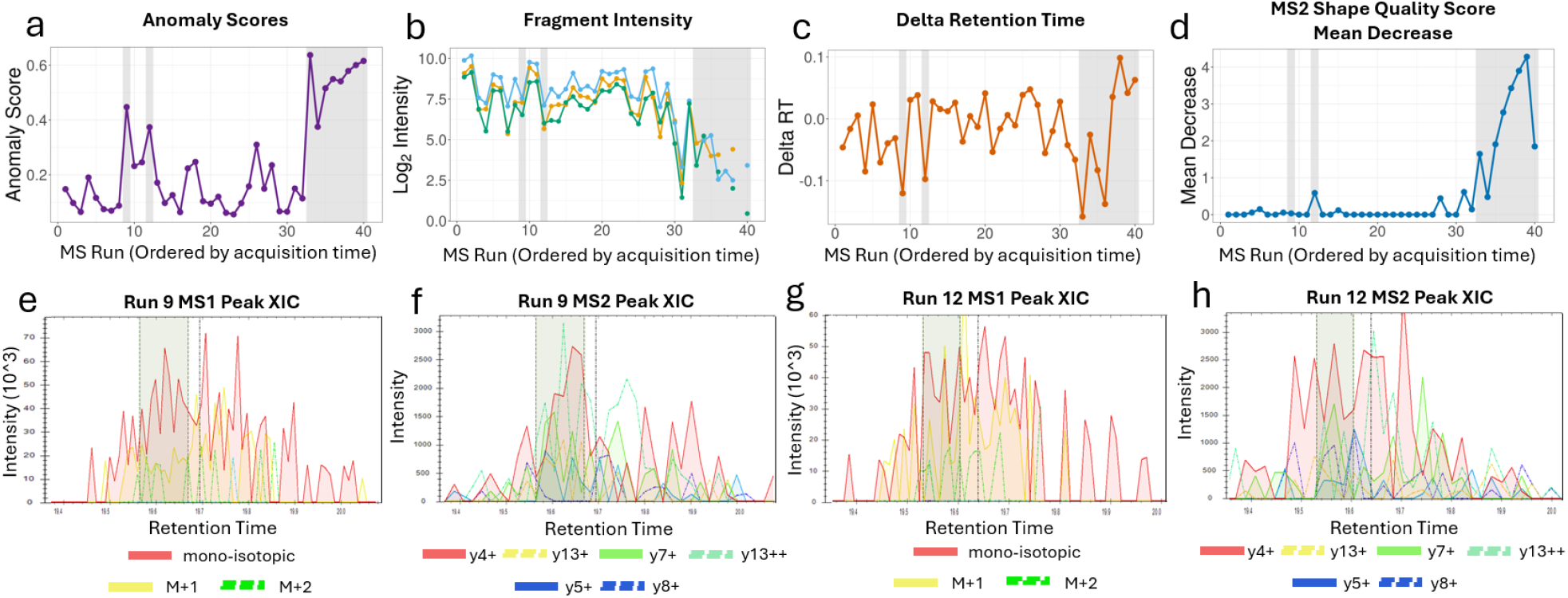
CSF benchmark, protein Q6UW01, precursor AAAGGPGGAAL-GEAPPGR.2: Anomaly scores captured drops in quality that were not seen from fragment intensities alone. Elevated anomaly scores (*>* .35) are highlighted in gray. **(a)** Precursor AAAGGPGGAALGEAPPGR.2 had elevated anomaly scores in runs 9, 12, and 33–40, indicating poor quality measurements. **(b)** The log_2_ intensities of the three fragments of the precursor were high in the anomalous runs, had no problematic values of other filtering criteria, and therefore were not captured by the preliminary filtering. The fragments are indicated with colors. **(c)** Deviations of the delta retention time metric indicated a problem with the measurements in runs 9 and 12, and in some but not all of the runs 33–40. **(d)** The engineered quality metric, negative CUSUM MS2 shape quality score, captured the longitudinal degradation drift in peak shape quality in runs 33–40. **(e)** The extracted ion chromatogram (XIC) for AAAGGPGGAALGEAPPGR.2 in run 9 at the MS1-level. Each colored line represents an isotopic peak of the precursor. The predicted retention time is marked by the black dotted line, while the observed retention time is highlighted in green. **(f)** The XIC in run 9 at the MS2-level. Each colored line represents a fragment ion of the precursor. The predicted retention time is marked by the black dotted line, while the observed retention time is highlighted in green. **(g)** Same as (e) for run 12. **(h)** Same as (f) for run 12.

Step 2 follows the workflow of base *MSstats*, and performs conservative univariate feature pre-filtering based on fragment loss, missing values, q-values of identification, and very small intensity values. It removes features and peak intensities that are obviously low quality, and ensures that the remaining low quality measurements are relatively rare. For example, in the case of Spectronaut this step removes fragments that lost a molecule (e.g., H_2_O, NH_3_, etc.) during the fragmentation process, have q-values above 0.01 (either on the protein group or elution group-level), or have missing values in more than half of the runs. The step also sets individual peak intensities to NA if they are below 1 or if Spectronaut marks them to be excluded from quantification. Full details of this step are given in Methods (**Section 4.2**). Finally, this step also includes log_2_ transformation and normalization as implemented in *MSstats*.

Step 3 replaces traditional feature selection with quality metric engineering and feature reweighting. *MSstats+* leverages peak quality metrics at the elution group and fragment group-levels, but does not use individual fragment quality information. This greatly improves the speed of the algorithm without sacrificing statistical accuracy (**Supplementary Section 2.2**). It begins by engineering additional quality metrics that capture longitudinal trends in quality of a precursor as a function of acquisition time. For example, positive and negative Cumulative Sums (CUSUMs) are transformations of a metric that are commonly used in longitudinal system suitability and quality control to capture drifts in mean or variation [43, 55].

Next, *MSstats+* summarizes all the original and the engineered metrics of a precursor in a run into a single anomaly score. This is done by means of isolation forest (iForest) [48], an unsupervised machine learning method that quantifies the deviation of a multivariate vector from its “typical” behavior. For each precursor, iForest builds multiple decision trees by recursively partitioning the runs in the quality metric space. It assumes that anomalous runs are relatively rare and easier to isolate in a binary tree structure, thus requiring fewer splits. Therefore, it defines the anomaly score of a run as a function of the average path length of the run over all the trees. Importantly, the anomaly score does not use the observed peak intensity as input and therefore is complementary to the intensity, avoiding overfitting, and can be calculated for precursors where the intensity is missing.

In Step 4, *MSstats+* performs two-step protein-level summarization, in a way similar, but not identical to base *MSstats* [30]. First, missing feature intensities are imputed using the Accelerated Failure Time (AFT) model in *MSstats*. As in *MSstats*, the imputation is performed for a fragment in a run where the protein has at least one other observed fragment (i.e., fragment intensities are not imputed in runs where the protein is entirely missing). Unlike *MSstats, MSstats+* views anomaly scores as proxies for the uncertainty associated with observed or imputed intensities. It incorporates the reciprocal of the anomaly scores into protein-level summarization as weights in a linear regression, such that highly anomalous intensities are downweighted. Unlike most existing protein-level summarization methods, the approach outputs not only an estimate of the protein-level summary in each run, but also the associated uncertainty quantified in terms of run-level variances, visualized as standard deviations (**Figure 1**). Anomalous measurements typically contribute to higher uncertainty of the protein-level summaries.

Finally, in Step 5, a linear mixed-effects model describes the systematic variation in protein-level summaries induced by the experimental design, as well as the random variation reflecting both the between-run biological and technological variation and the within-run unequal variance of protein-level summaries. To account for the unequal variance, the parameters of the model are estimated by weighted maximum likelihood (or weighted restricted maximum likelihood depending on the design, following *MSstats*). Finally, similarly to *MSstats*, comparisons between conditions of interest are specified as model-based contrasts.

### 2.2. Open-source implementation of *MSstats+*

*MSstats+* is implemented as part of the family of open-source R/Bioconductor packages *MSstats* [30, 52–5*4]. The implementation is fully integrated with existing MSstats* features such as data processing (e.g., converters to upstream data processing tools, pre-filtering and normalization, metadata structures describing the experimental design), statistical model support for complex experimental designs (e.g., repeated measure designs, advanced comparisons between conditions and covariate adjustment), and a variety of plotting functions. To help evaluate the extent of anomalous measurements in a dataset, *MSstats+* implements inside the converter package *MSstatsConvert* a new function `CheckDataHealth`, which summarizes precursor-level anomaly scores with Pearson’s moment coefficient of skewness.

Since anomaly scores are calculated at the precursor level, efficiency and scalability are critical. *MSstats+* includes a custom C++ implementation of the isolation forest using the `Rcpp` package [56], significantly improving speed over native R implementations. Additionally, *MSstats+* supports multi-core parallelization, distributing precursor-level isolation forests across threads. These optimizations support scalability to large experimental datasets (see **Supplementary Fig. 4** for comparisons between the R and C++ implementations and **Supplementary Fig. 5** for a scalability analysis). Beyond iForest, *MSstats+* supports large-scale experiments through *MSstatsBig*, which leverages out-of-memory data handling and filtering prior to protein summarization and differential analysis [50].

**Figure 4.**
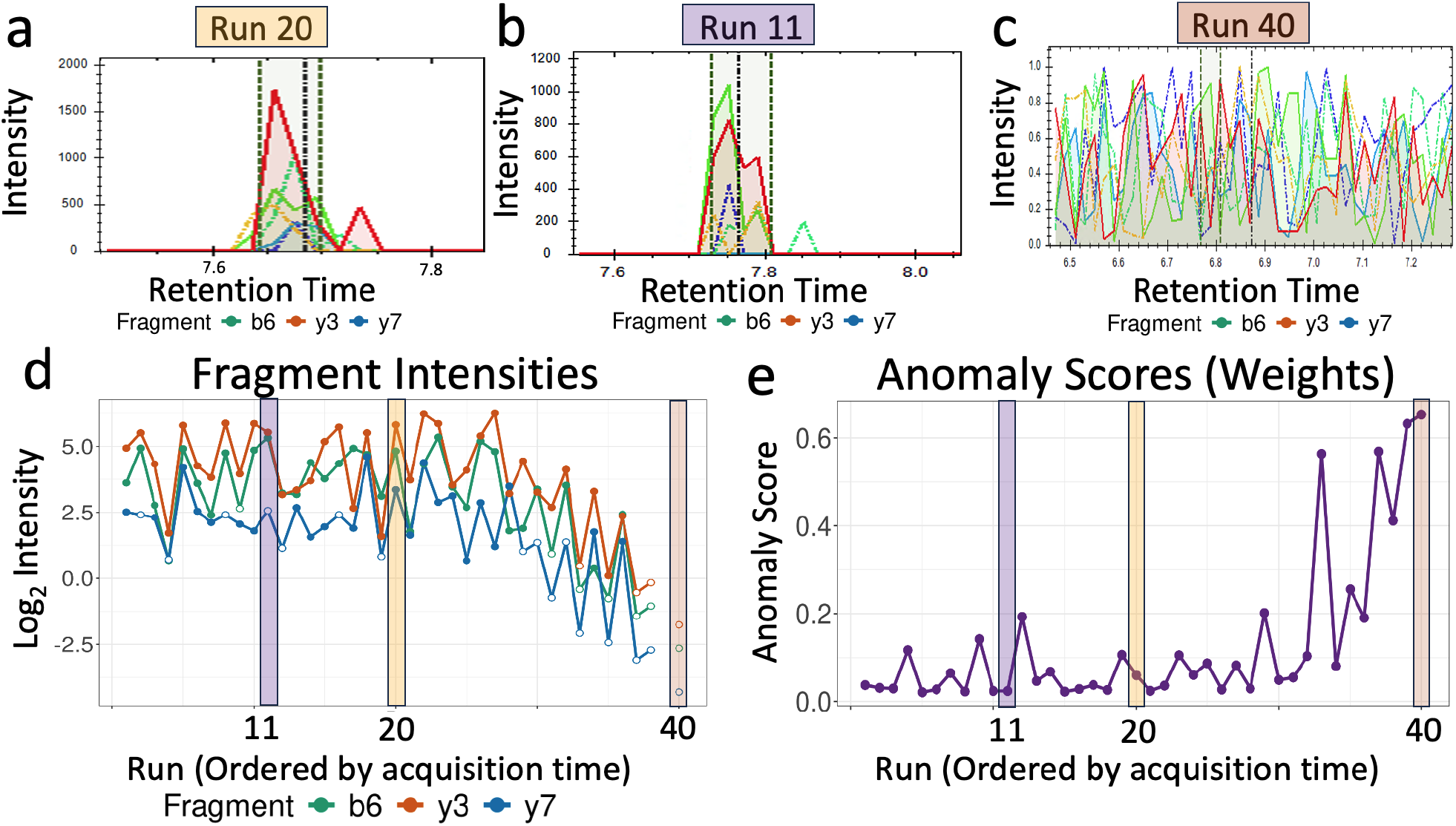
K562 benchmark, protein Q9UII2, precursor EAGGAFGK.1: anomaly scores reflected confidence in fragment-level imputation. **(a)** Fragment peaks in run 11 (b6 - solid green, y3 - solid red, y7 - solid blue, y6 - dotted yellow, b7 - dotted green, y5 - dotted blue). The precursor had no missing intensities in run 20. **(b)** Fragment peaks in run 11. Colors as in (a). The peaks for b6 and y3 fragments were well-defined. The y7 fragment was missing and was marked for imputation by *MSstats+*. **(c)** Fragment peaks in run 40. Colors as in (a). The run had no clearly defined precursor peak and had noisy fragments. All the fragment in this run were marked for imputation by *MSstats+*. **(d)** The y6, b7, and y5 fragments were pre-filtered by *MSstats+*, therefore the precursor was quantified by b6, y3, and y7 fragments. As expected from the experimental design, the intensities of the fragments decreased in later runs. While most of the peak intensities were fully observed (solid colored points) such as in run 20 (yellow band), others were imputed (hollow white points), such as in runs 11 and 40 (purple and brown bands). **(e)** The anomaly scores for the precursor drifted upwards in later runs. The inverse of these scores were used to weight the observed and the measurements in (a). The imputed value in run 11 had a low anomaly score and a high weight, while the imputed values in run 40 had a high anomaly score and a low weight.

**Figure 5.**
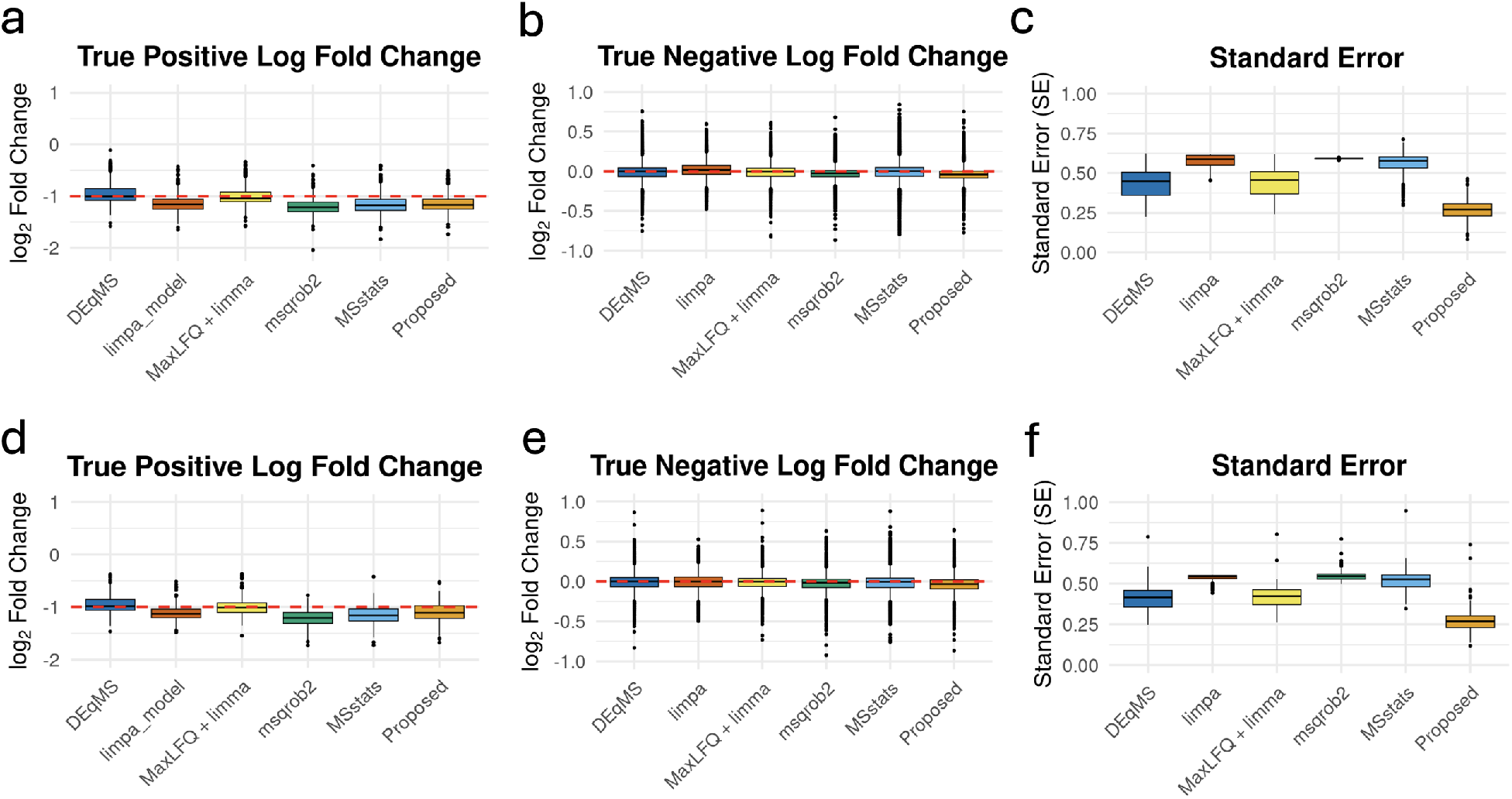
K562 and CSF benchmarks: The proposed approach increased experimental efficiency without introducing bias. (**a**) In K562 benchmark, most statistical methods accurately estimated the log_2_ fold changes of the true differentially abundant proteins. (**b**) In K562 benchmark, all the statistical methods accurately estimated the log_2_ fold changes of the true non-differentially abundant proteins. (**c**) In K562 benchmark, *MSstats+* produced consistently lower estimates of standard errors. (**d-f**) As in (**a**-**c**), the CSF benchmark.

### 2.5. On benchmark experiments with known anomalies, *MSstats+* outperformed existing statistical methods

#### 2.3.1. Evaluation strategy

We evaluated the proposed approach on two benchmark experiments, specifically designed to contain both known ground truth of differential abundance and known anomalous measurements. The first experiment profiled K562 cell lysate, and the second profiled pooled cerebrospinal fluid (CSF) from human subjects. The CSF benchmark had greater dynamic range and included low abundant analytes, making the sample more challenging to measure and increasing the technical variability. This made the CSF benchmark more representative of a real-world proteomic investigation and more challenging for statistical analyses (in comparison to the K562 benchmark).

**Figure 2a** illustrates the dilution scheme used for both the K562 and CSF benchmark experiments. In both experiments, we manually introduced two conditions by varying the amount of biological material injected per run, mimicking a biological group comparison with a known log_2_ fold change of 1. Anomalous runs were introduced by gradually injecting lower amounts of material, mimicking the effects of instrumental drift and degradation in measurement quality over time. Reduced material increases the likelihood of unstable peak shapes, retention time deviations, and other indicators of poor quantification, however, it does not mean that an anomalous run is simply less intense. In real experiments, anomalies are not defined by intensity, but by deviations in quality metrics. Finally, we assume that instrumental drift occurs on an individual protein basis, such that it cannot be corrected by global normalization or by censoring entire runs.

Data were acquired using an Orbitrap Astral Mass Spectrometer coupled to a Thermo Scientific Vanquish Neo UHPLC with DIA acquisition and a total of 40 runs in each experiment. The resulting datasets were processed using Spectronaut v19. In addition to precursor and fragment intensities, the report included quality metrics such as delta retention time, peak shape score on the MS1 and MS2 levels (derived metrics from Spectronaut integrating several aspects of signal quality), and acquisition time stamp. These metrics were reported per-precursor, such that all the fragments of a precursor had the same values of the metrics. To mimic a common situation where most proteins do not change in abundance between conditions, we randomly shuffled condition labels of the runs for 90% of the quantified proteins, eliminating the differences between conditions, while preserving the true run labels and fold changes for the remaining 10% of the quantified proteins. Further details are provided in **Supplementary Table 6** and the Methods (**Section 4.1**).

We compare *MSstats+* to five of the most popular statistical methods used to analyze MS proteomics experiments: base *MSstats, msqrob2, DEqMS, limpa*, and *limma*. All of the methods except for *limma* include their own protein-level summarization functionalities. For *limma* we leverage *MaxLFQ* summarization implemented in the *iq* R package [57].

#### 2.3.2. One protein example

**Figure 2** illustrates *MSstats+* in the case of protein A4D126 from the K562 benchmark and in particular its precursor ALAEDQINSK.2, both of which were randomly selected. The intensity of the precursor in run 40 was unremarkable (its log_2_ intensity of 2.41 fell in the 22.5^th^ percentile of precursor intensities), and we would not be able to diagnose quality issues with that run from the intensity value alone (**Figure 2b**). Beyond intensity, the standard filtering cutoffs (such as q-value and the exclusion from quantification filter) did not mark this measurement for removal. However, the precursor had only one detected fragment ion, and showed a larger than typical drop in MS2 shape quality score. For this precursor, the discrepancy was captured by the temporal transformation of the “MS2 Shape Quality Score” metric in Spectronaut (negative CUSUM) (**Figure 2c**) and resulted in a high overall anomaly score of approximately 0.6 (**Figure 2d**). Additional quality metrics for this precursor can be found in **Supplementary Fig. 6**.

**Figure 6.**
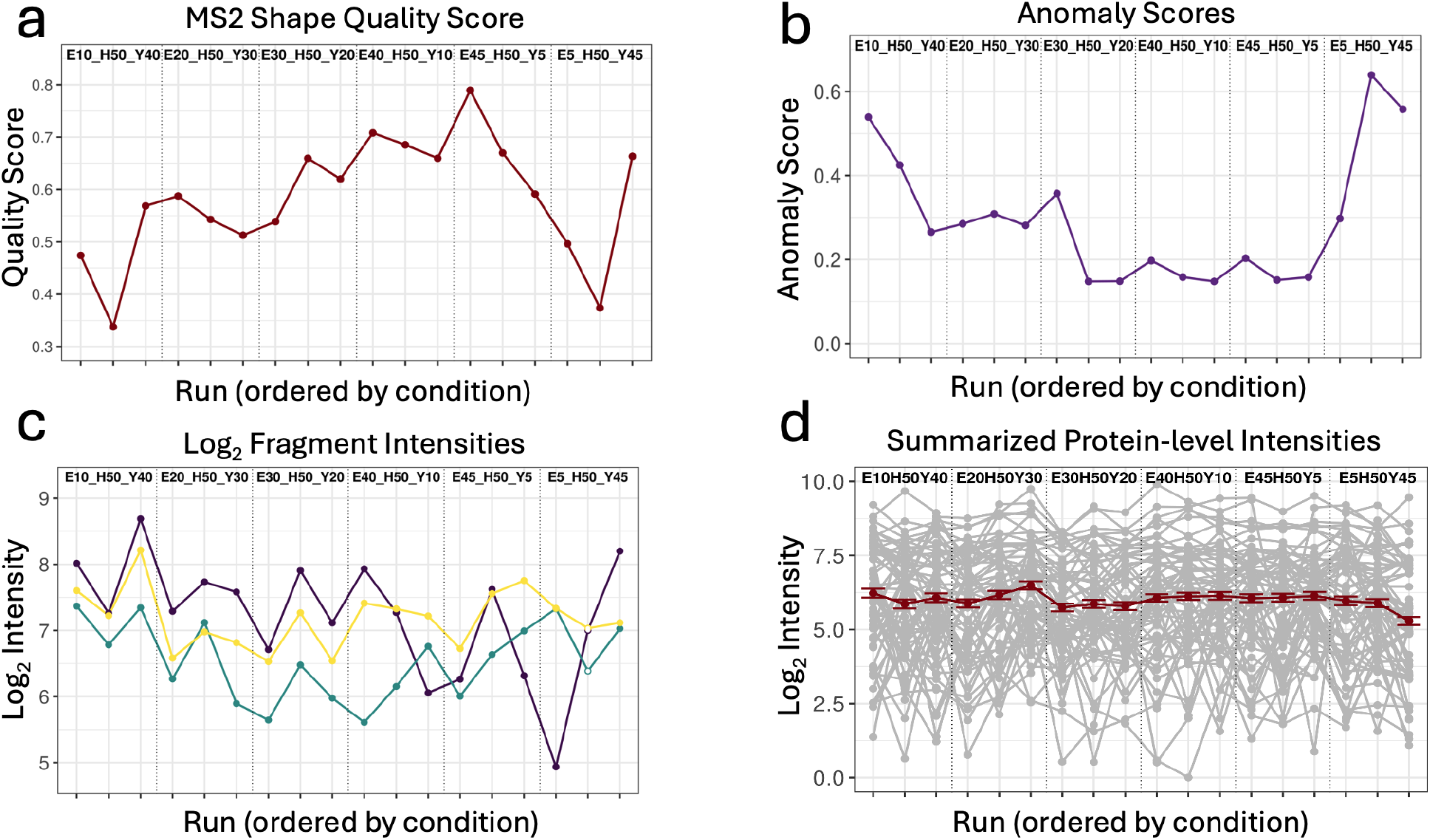
Mixture of proteomes, human protein O15091: *MSstats+* downweighted noisy measurements that were interpreted as biological variation by other statistical methods. **(a)** Precursor SRPPFDVVIDGLNVAK.3 had a lower than typical shape quality score at the MS2-level in conditions E5 H50 Y45 and E10 H50 Y40, indicating measurement issues. **(b)** Anomaly scores for precursor SRPPFDVVIDGLNVAK.3 captured these issues, and were higher in the problematic runs. **(c)** SRPPFDVVIDGLNVAK.3 intensity values indicate some increased noise in condition E5 H50 Y45. In particular, the b7 ion (colored purple) exhibited increased variability across all replicates in this condition. In comparison, the intensity values in condition E10 H50 Y40 did not exhibit increased noise, even though the anomaly scores for these runs were high, indicating the model detected drops in quality that were not discernible from intensity values alone. **(d)** *MSstats+* summarized the fragments into a single protein-level abundance per run. The problematic runs had higher associated uncertainty, and were downweighted in the subsequent statistical analysis. The red line represents the summarized protein-level intensities and the error bars show the standard deviation of the run-level estimates.

All the precursors of A4D126 were analyzed in the same way. The anomaly scores of the later runs were higher for all precursors (**Figure 2e**), reflecting the experimental design that intentionally introduced anomalies in the later runs. Beyond this longitudinal trend, individual anomalous precursors were also observed. For example, in Run 7 the shape quality score at the MS2-level showed a substantial drop, resulting in a higher than average anomaly score for that run. This illustrates the ability of *MSstats+* to detect both longitudinal trends in quality degradation and non-systematic quality drops.

To aggregate the fragment intensities in each run into protein-level summaries, *MSstats+* used the inverse of the anomaly scores as weights, such that intensities with higher anomaly scores contributed less to the summary (**Figure 2f**). As a result, later runs were associated with both lower values of protein summaries, and higher uncertainty in the summaries (represented by error bars showing the standard deviation of the summaries).

The table in **Figure 2e** contrasts the differential analyses of A4D126 between *MSstats+* and *MSstats. MSstats+* estimated a more accurate log_2_ fold change (true value: -1), reduced the associated standard error by 36%, and identified the protein as differentially abundant. We performed differential analysis of this protein with other statistical methods (**Supplementary Table 2**), and *MSstats+* was the only method able to detect this protein as differentially abundant.

Additional examples demonstrating that *MSstats+* is consistent across proteins can be found in **Supplementary Fig. 7**–**Supplementary Fig. 13** and **Supplementary Table 3**.

**Figure 7.**
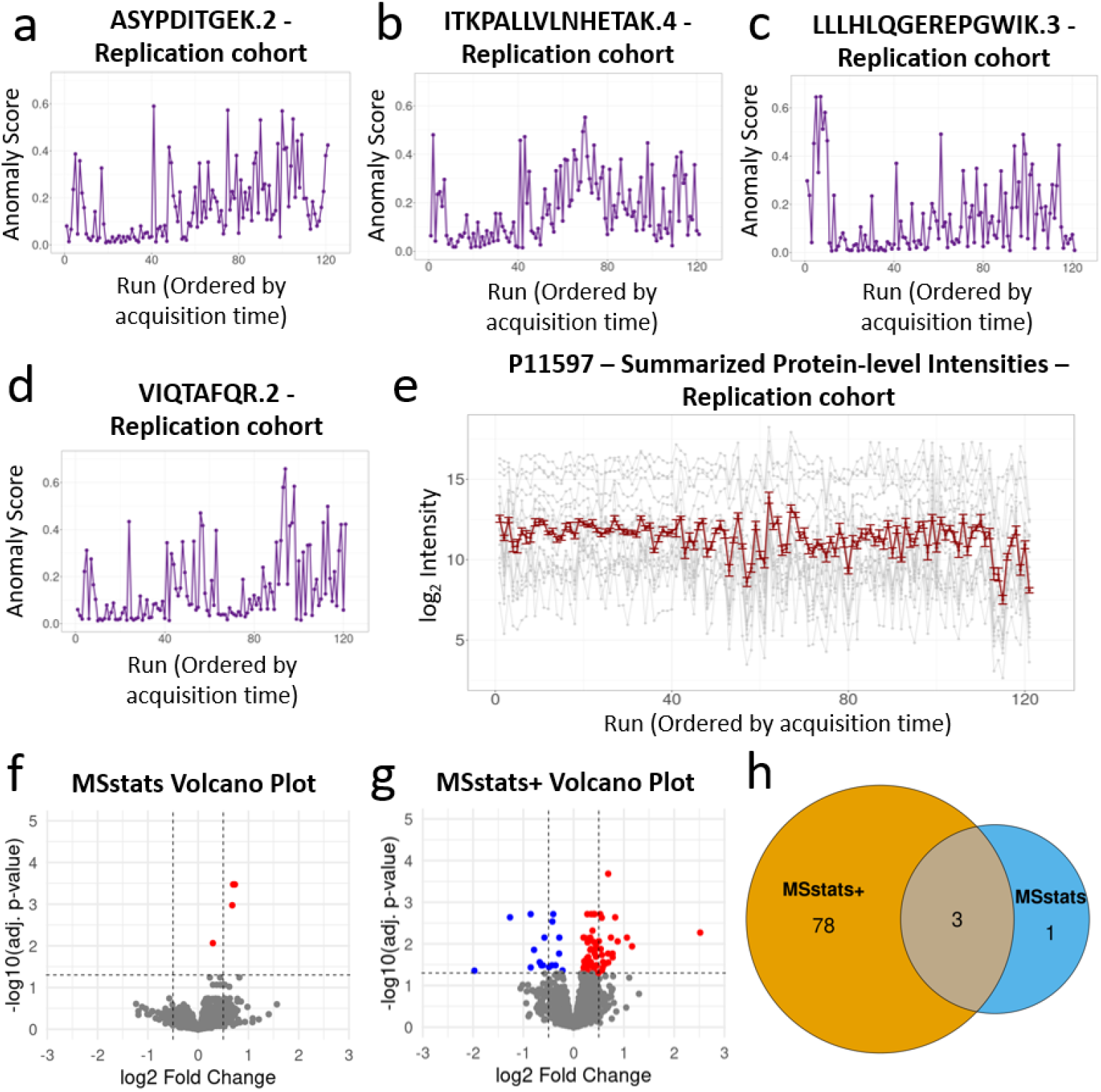
Clinical study: *MSstats+* corrected for poorly quantified measurements, and identified more differentially abundant proteins compared to *MSstats*. (**a)** Anomaly scores for precursor ASYPDITGEK.2 in protein P11597. **(b)** Anomaly scores for precursor ITKPALLVLNHETAK.4 in protein P11597. **(c)** Anomaly scores for precursor LLLHLQGEREPGWIK.3 in protein P11597. **(d)** Anomaly scores for precursor VIQTAFQR.2 in protein P11597. **(e)** Protein-level summary of protein P11597. Grey lines show fragment intensities, the dark red line represents the protein-level summary. Red error bars are standard deviations of the run summaries. **(f)** Differential analysis results from *MSstats* shown as a volcano plot. Up-regulated proteins are shown in red, down-regulated in blue. *MSstats* identified four proteins as up-regulated. **(g)** Same as (f) for *MSstats+*. By downweighting the problematic runs, *MSstats+* reduced the standard errors and identified more differentially abundant proteins (both up- and down-regulated). **(h)** Overlap of differentially abundant proteins detected by *MSstats+* and *MSstats*, with 3 shared, 1 unique to *MSstats*, and 78 unique to *MSstats+*.

#### 2.3.3. *MSstats+* detected anomalies that were not seen from peak intensities alone

The K562 and CSF benchmarks intentionally introduced low quality measurements. To evaluate the ability of *MSstats+* to detect anomalous measurements in these datasets, we calculated Pearson’s moment coefficient of skewness [58] for each precursor. For each precursor, most runs are expected to receive low anomaly scores with a few high-scoring outliers (i.e., a right-skewed distribution). Pearson’s moment coefficient of skewness quantifies this trend, with a positive coefficient indicating a skew and a coefficient near 0 indicating no skew. We visualize the coefficients of skewness calculated for all the precursors in each benchmark with a histogram (**Supplementary Fig. 14a-b**). The histograms of the coefficients were shifted away from zero with the average close to 1. This indicates that the algorithm was able to detect the presence of anomalous low quality measurements.

Intentionally introduced anomalies were strongly associated with higher anomaly scores in both benchmarks. As shown in **Supplementary Fig. 15**, the distributions of the anomaly scores shifted toward larger values in the anomalous measurements, indicating that *MSstats+* detected the anomalous measurements correctly.

Importantly, the anomaly scores were not proxies of the fragment intensities. As shown in **Supplementary Fig. 16**a–b, the anomaly scores had only a weak correlation with log_2_ intensities in both benchmarks. Although low-intensity measurements were somewhat more likely to have elevated anomaly scores, many low-abundant measurements remained highquality, and conversely, some high-intensity measurements had anomalous behavior. In other words, the anomaly scores captured quality information beyond simple signal strength, supporting their use in downstream statistical analysis.

**Figure 3** illustrates a weak association between fragment intensity and quality score for the precursor AAAGGPGGAALGEAPPGR.2 of the protein Q6UW01 in the CSF benchmark. The precursor had elevated anomaly scores in runs 9, 12, and 33–40 (**Figure 3a**). However, the log_2_ intensities of the three fragments of the precursor as well as other criteria used for filtering were within the typical range (**Figure 3b**), and were not captured by preliminary filtering. Deviations in the delta retention time quality metric indicated an underlying problem with the measurements in runs 9 and 12 (**Figure 3c**). The additional engineered quality metric, negative CUSUM MS2 shape quality score, captured another dimension, the longitudinal degradation drift in MS2 shape quality in runs 33–40, resulting in high anomaly scores for these runs (**Figure 3d**). **Supplementary Fig. 17** highlights additional quality metrics for this precursor. In run 9, the MS1 shape quality score showed a significant drop, while in run 12 there was a drop in the MS2 shape quality score. These deviations provide additional evidence of measurement issues in these samples, even though the precursor intensities were reasonable.

To validate the observations from the quality metrics, we manually inspected runs 9 and 12 using the extracted ion chromatograms (XIC) at the MS1and MS2-levels (**Figure 3e-h**). In run 9, both MS1 and MS2 XIC traces showed poor alignment and increased noise across the retention time window. In run 12, the MS2-level XIC was particularly noisy, and the y4 ion (colored red) showed multiple peaks near the predicted retention time. These manual inspections were consistent with the anomalies identified by *MSstats+*.

#### 2.3.4. Anomaly scores naturally reflected confidence in imputed missing values

*MSstats+* leverages how Spectronaut and other spectral processing tools report missing values to improve imputation. Spectronaut rarely reports entirely missing values (i.e., “NA” peak intensities). Instead, when a fragment peak cannot be reliably located, it integrates the noise in the expected region of the peak and reports low intensities (less than 1). *MSstats+* treats these low fragment intensities as missing (“NA”) but retains their underlying peak quality metrics as informative of the missingness. If a precursor is partially observed and only some fragment ions are missing, the missing fragments have precursor-level scores. If a precursor is entirely missing in a run, its quality metrics reflect the integrated noise and are typically low.

**Figure 4** illustrates the ability of *MSstats+* to handle missing values, in the case of precursor EAGGAFGK.1 of protein Q9UII2 in the K562 benchmark. In run 20 (**Figure 4a**), all fragments of the precursor were observed, the spectral peaks were well-defined, and the anomaly scores were low. Run 11 (**Figure 4b**) displayed clean spectral peaks for the b6 and y3 fragments, but fragment y7 was missing. The missing intensity value was imputed by *MSstats+* based on the value of the observed fragments in this run, and based on the values of the missing fragment in the runs where it was observed. Despite the missing value, the precursor had high quality metrics. Therefore, the anomaly score of the imputed value was low, indicating that the imputation was done based on high information content. In contrast, run 40 (**Figure 4c**) had no clearly defined precursor peak and had noisy fragments. *MSstats+* imputed all the fragment values of this precursor in this run based on the observed fragments of other precursors of the same protein in this run, and based on the values of the missing fragments in the runs where they were observed (**Figure 4d**). However, the anomaly scores of this precursor indicated low information content (**Figure 4e**). As a result, this precursor was downweighted by protein summarization, and the summary had higher associated uncertainty.

Additional examples illustrating the treatment of missing values under varying spectral conditions are in **Supplementary Fig. 18** and **Supplementary Fig. 19. Supplementary Fig. 20** illustrates, at the experiment-wide level for both benchmarks, that the anomaly scores of the imputed intensities span the entire range of values. In other words, *MSstats+* distinguished informative and non-informative missing values by leveraging their underlying quality metrics, rather than indiscriminately downweighting all the imputed values.

#### 2.3.5. *MSstats+* improved detection of differentially abundant proteins

All statistical methods evaluated in this manuscript accurately estimated log_2_ fold changes in the proteins in the K562 and CSF benchmarks, with only minor differences between methods (**Figure 5a-b** and **Figure 5 d-e**). However, substantial differences emerged in the estimation of standard error (**Figure 5c** and **Figure 5f**). *MSstats+* consistently produced the lowest standard errors across both benchmarks. Both *DEqMS* and *limma* produced the second lowest standard errors (after *MSstats+*), due to their use of empirical Bayes methods that shrink variance estimates across proteins.

Due to its ability to downweight runs with low quality quantifications and to reduce standard errors, *MSstats+* substantially increased the accuracy of detecting differentially abundant proteins. **Table 1** summarizes the performance across all the statistical methods. *MSstats+* correctly identified 91.4% and 86.9% of true positives in the K562 and CSF datasets, respectively, without introducing false positives. The lower proportion of true positives in the CSF benchmark reflected the higher complexity of the CSF samples. Importantly, improved performance was achieved without double-dipping into the data, avoiding inflating the false positive rate to increase sensitivity. In contrast, all other methods failed to detect true positives due to inflated standard errors.

**Table 1.** Controlled experiments with known ground truth: *MSstats+* improved detection of differential abundance. Metrics are true positive rate (TPR), and positive predictive value (PPV). *MSstats+* was the only method able to detect true positives in the K562 and CSF benchmarks with intentionally introduced anomalies. It maximized the positive predictive value in the mixture of proteomes.

| Method | Dataset 1:<br>K562 benchmark |  | Dataset 2:<br>CSF benchmark |  | Dataset 3:<br>Mixture of proteomes |  |
| --- | --- | --- | --- | --- | --- | --- |
|  | TPR | PPV | TPR | PPV | TPR | PPV |
| MSstats+ | 0.914 | 1.000 | 0.869 | 1.000 | 0.794 | 0.684 |
| MSstats | 0.000 | NA | 0.000 | NA | 0.826 | 0.616 |
| limpa | 0.000 | NA | 0.000 | NA | 0.845 | 0.659 |
| MaxLFQ + limma | 0.000 | NA | 0.000 | NA | 0.820 | .589 |
| msqrob2 | 0.000 | NA | 0.000 | NA | 0.843 | 0.626 |
| DEqMS | 0.000 | NA | 0.000 | NA | 0.781 | 0.628 |

This result is expected given the design of the benchmark studies. The manually introduced instrumental drift was designed to mimic unreliable measurements that increase technical variance without biasing the fold change. Existing statistical methods do not downweight unreliable measurements and interpret this drift as biological variability, inflating standard errors to the point that no true positives can be detected. In comparison, *MSstats+* explicitly downweights poorly quantified measurements, controlling the standard error, and preserving statistical power to detect differentially abundant proteins.

### 2.4. In the controlled mixture of proteomes, *MSstats+* improved the positive predictive value of differential abundance

We further evaluated *MSstats+* on a previously published controlled mixture of proteomes, designed to represent a proteomic experiment with known ground truth and no deliberately introduced anomalies [49]. Tryptic peptides from three different species, human, yeast, and *Escherichia coli* (*E. coli*), were combined, such that the relative abundance of yeast and *E. coli* proteomes varied in six different ratios, and the human proteome was kept constant. More details on the data processing, number of quantified proteins and the experimental design can be found in Methods (**Section 4.1**).

This experiment featured a relatively low number of replicate runs and a highly optimized Astral DIA workflow. The resulting dataset was markedly clean, with 90% of the quantified proteins showing CVs below 20% and a low number of missing values [49]. Consistent with this observation, the experiment showed a distribution of Pearson’s moment coefficient of skewness fluctuating around 0 (**Supplementary Fig. 14c**), indicating a lower prevalence of poor quality measurements. *MSstats+* is designed for large scale and high complexity datasets of heterogeneous quality and is expected to provide the greatest improvement over standard approaches when poor quality measurements are present. However, even in cleaner experiments, the method can still outperform standard approaches and does not introduce additional bias.

We compared the performance of differential analysis by *MSstats+* to the same statistical methods as in the other benchmarks. The six mixtures were compared to each other in pairwise comparisons. Since human proteins were mixed at a constant ratio, they served as true negatives in all comparisons, while yeast and *E. coli* proteins were treated as true positives. Despite the relatively rare prevalence of anomalous measurements, *MSstats+* performed favorably (**Table 1**). It achieved the highest positive predictive value (PPV of 68.4%) and a comparable true positive rate (TPR of 79.4%), without inflating false discoveries. The other statistical methods had lower positive predictive values and more false positive differentially abundant proteins. A summary of the differences between the estimated and ground truth fold changes, as well as a comparison of the standard errors across all statistical methods evaluated can be found in **Supplementary Fig. 21**.

**Figure 6** illustrates one potential source of false positives in the case of the human protein O15091. The protein did not have a true change in abundance in any of the conditions. The precursor SRPPFDVVIDGLNVAK.3 had high MS2 shape quality score in the majority of runs indicating good quality measurements, but dropped to lower values in both condition E5 H50 Y45 and E10 H50 Y40 indicating measurement issues (**Figure 6a**). Anomaly scores for the precursor captured these issues and were higher in the problematic runs (**Figure 6b**). The problematic runs shifted the fragment intensities, introducing a bias (**Figure 6c**). *MSstats+* summarized the fragments into a single protein-level abundance per run (**Figure 6d**). Since the problematic runs had higher associated uncertainty, they were downweighted in the subsequent statistical analysis. *MSstats+* was able to incorporate the poor quality measurements into differential analysis, and lowered the estimated log_2_ fold change while retaining a high standard error. Across all 15 comparisons for O15091, *MSstats+* detected only one false positive, while the other statistical methods each detected five or more (**Supplementary Table 4**).

### 2.5. In a clinical cohort *MSstats+* increased the sensitivity of differential analysis

We evaluated the performance of *MSstats+* in a clinical proteomic cohort from the Tauriel and Lauriet Phase II clinical trials, which studied the safety and efficacy of Semorinemab in Alzheimer’s disease [51]. More than 250 cerebrospinal fluid (CSF) samples were collected at baseline and follow-up time points (49, 61, or 73 weeks). Data were acquired with DIA on an Orbitrap Exploris 480 Mass Spectrometer, processed with Spectronaut v19, and identified over 3000 proteins. More details on the experimental design are available in **Supplementary Table 6** and the Methods (**Section 4.1**).

For differential analysis, we compared subjects with mild versus more advanced cognitive impairment at all time points, as defined by the Clinical Dementia Rating Scale Sum of Boxes (CDR-SB) score cutoff of 4.5. This cutoff was based on prior clinical studies [59] and by the empirical distribution of participant CDR-SB scores to ensure broadly balanced groups (**Supplementary Fig. 22**).

The study had a large number of runs, high sample complexity, and required complex sample preparation steps such that deviations from optimal performance were unavoidable in at least some of the measurements. Indeed, the experiment showed a distribution of Pearson’s moment coefficient of skewness near 1 (**Supplementary Fig. 14**d), indicating that *MSstats+* was able to detect poorly quantified runs in most precursors.

**Figure 7**a-d highlights the anomaly scores calculated by *MSstats+* for four precursors of protein P11597 (CETP). Across all precursors the first few runs showed elevated anomal scores (particularly in runs 5 and 7), suggesting unreliable quantification. After these early runs, anomaly scores were generally lower until run 41, where they rose sharply. After run 41, precursors displayed highly variable anomaly scores, with large jumps and occasional low values until the end of acquisition. These trends carried through to the protein-level summarization in **Figure 7**e. The variance estimates (error bars) were elevated in the first few runs, decreased until run 40, and then increased markedly after run 41. These observations can be further validated by the log_2_ intensity measurements of P11597. Apart from the first few runs, intensities prior to run 41 showed low variability. From run 41 onward, the variability increased substantially. This trend follows the anomaly score patterns in **Figure 7a-d**, despite intensities not being used to estimate the anomaly scores.

Finally, we contrast the differential analysis for P11597 between *MSstats+* and *MSstats* (**Supplementary Table 5**). *MSstats+* downweighted the poorly-quantified runs using the variances in **Figure 7e**. Compared to *MSstats*, this resulted in a similar fold change estimate, but a reduced standard error and a statistically significant adjusted p-value (*α <* 0.05). Identifying P11597 as differentially abundant aligns with prior studies linking CETP to Alzheimer’s disease pathology [60, 61].

**Figure 7f-h** compares the differentially abundant proteins detected by *MSstats+* and *MSstats*. By downweighting the problematic runs (without entirely excluding them from the run-level summarization), *MSstats+* reduced the standard errors and identified more differentially abundant proteins. Three of the proteins were shared with *MSstats*, and one was unique to *MSstats*. The protein unique to *MSstats* was Q14767 (LTBP2), which had the lowest fold change among all proteins identified as differentially abundant by *MSstats* (**Figure 7f**). In contrast, *MSstats+* downweighted the quantitative measurements for Q14767, resulting in an increased standard error and higher p-value. For the other three proteins identified by *MSstats, MSstats+* considered the measurements more reliable, leading to similar fold changes and standard errors across both methods.

Although the true differentially abundant proteins in this study are unknown, the set of proteins uniquely identified by *MSstats+* includes several known to be associated with Alzheimer’s disease pathology, including CETP, NTRK1 [62], SERPINE2 [63], SNCB[64], EFHD2 [65], and PIN1 [66]. Full differential analysis results, and comparisons with *MSstats*, are provided in **Supplementary Data 1**.

## 3. Discussion

We presented *MSstats+*, a comprehensive statistical workflow for incorporating quality metrics into differential analysis. *MSstats+* supports large-scale studies where manual quality checks are impractical or not possible at all. *MSstats+* downweights anomalous measurements based on a multivariate pattern of their quality metrics that are not easy to detect by simple univariate filtering. Since quality metrics contain information that is complementary to the information in feature intensities, *MSstats+* avoids double-dipping and protects against overfitting. *MSstats+* offers a natural treatment of imputed missing values, as the quality metrics characterize the quality of the information content used for the imputation. As a result of the methodological advances, *MSstats+* improves detection of differentially abundant proteins. In our evaluations *MSstats+* improved the detection of differentially abundant proteins, while controlling FDR, compared to existing statistical analysis methods. In particular, *MSstats+* demonstrated strong performance in the presence of precursor-level instrumental drift, a situation that is easily overlooked by standard pipelines.

*MSstats+* detects poor-quality measurements at the precursor-level, despite relying on fragment-level quantification for summarization. The model could theoretically be applied at the fragment-level by leveraging individual fragment peak quality metrics, potentially capturing quality drops that may be missed at the precursor-level. However, in our tests, this approach did not improve accuracy and significantly increased computational cost due to the large number of isolation forest models required. Future work could explore precursor- versus fragment-level modeling in more detail, potentially applying fragment-level models only when warranted.

In the analyses presented in this manuscript, *MSstats+* partially relied on Spectronaut’s internal calculation for MS1and MS2-level peak shape quality. Spectronaut’s shape quality score integrates several aspects of signal quality, including mass-offset, XIC-shape, and the isotopic distribution correlation, into a single metric ranging from -1 to 1. By using these integrated scores, we avoided relying on an extensive list of separate quality metrics, while still capturing multiple dimensions of measurement quality. This approach provided a balance between comprehensiveness (i.e., incorporating many aspects of quality) and interpretability. However, relying on aggregated metrics may cause *MSstats+* to overlook aspects of peak quality not fully captured by the metric. Future work, both using Spectronaut and other tools, should compare the use of aggregated metrics (e.g., peak quality score) against a broader set of individual metrics.

Although the examples in this manuscript use Spectronaut as a data processing tool, *MSstats+* is not tool-specific and can be applied to any data processing tool that exports peak identification and quantification quality metrics. The methods in *MSstats+* are implemented directly in the *MSstats* converter for Spectronaut, which allows the user to simply specify which peak quality metrics to leverage in the model. For other tools, while the model itself is universal, manual formatting is currently required to put the data in the correct format for the model. An example of how to do this is provided in the publicly available *MSstats+* vignette. Future implementation work will involve integrating the model directly into all DIA converters in *MSstats*.

A natural question is how *MSstats+* extends to cases with highly variable samples, such as those at very low abundances (i.e., near the limit of detection) or generated from small amounts of biological material (e.g., single-cell proteomics). By design, the anomaly detec-tion model in *MSstats+* identifies measurements that deviate from the bulk of observations. Thus, in experiments with high variability, where measurements are spread across a wide range, the method highlights only the most extreme outliers, whereas in less variable experiments it flags smaller deviations. This property suggests that the approach should scale well to more variable settings, including single-cell studies. A future direction is to benchmark and refine the method for such applications.

Although this work focuses on DIA acquisition, the method can, in principle, be extended to quantitative experiments with other acquisition types. This includes label-free data-dependent acquisition (DDA), as well as experiments using tandem mass tags (TMT) labeling. Likewise, the approach could be adapted to studies interested in biological questions beyond changes in protein abundance, such as changes in post-translational modifications (PTMs) or protein structural changes (using methods such as limited proteolysis (LiP)). Potential challenges with these extensions include limited availability of spectral features (both due to MS1-level quantification and peptide-level analysis), different spectral peak quality metrics (on the MS1-level), and noisier peptide-level measurements (in the case of PTM or LiP). Future work should focus on extending the proposed method in these directions.

The method is implemented in the family of open-source R/Bioconductor *MSstats* packages that support a variety of complex experimental designs, allowing users to easily integrate and adapt it into their own workflows. Overall, the statistical methodology and the practical implementation of *MSstats+* make it poised to become a critical tool for quantitative mass spectrometry-based proteomic investigations.

## 4. Methods

### 4.1. Data overview and availability

#### K562 and CSF benchmarks

##### Sample preparation

For K562 proteome profiling, neat K562 samples were prepared by re-suspending 100 ug of K562 digest (Promega, V6951) in 100 ul of 0.1% Formic Acid, 0.01% n-dodecyl *β*-D-maltoside detergent, and 0.006% 10X iRT (Biognosys). These neat K562 samples were then further diluted with 0.1% Formic Acid, 0.01% n-dodecyl *β*-D-maltoside detergent, and 0.006% 10X iRT (Biognosys) to achieve dilutions of 1:2, 1:4, 1:8, 1:16, 1:32, and 1:64. For CSF proteome profiling, 25 µL of CSF from hydrocephalic donors (referred to as neat CSF) underwent reduction, alkylation, and digestion using trypsin for 3 hours, followed by peptide clean-up. Peptide quantification was performed using the Pierce Quantitative Fluorometric Peptide Assay (Thermo Scientific, 23290). The peptides were then dried in a SpeedVac (SP Scientific). Neat CSF peptides at a concentration of 0.1 µg/µL were further diluted to match the dilutions used for K562 samples.

##### Orbitrap Astral LC–MS/MS analysis

The analysis was conducted on an Orbitrap Astral Mass Spectrometer coupled to a Thermo Scientific Vanquish Neo UHPLC. Neat K562 and CSF peptides were separated using a reversed-phase C18 Ion Opticks 25 cm x 75 *µ*m ID, 1.7 *µ*m column on a Thermo Scientific Vanquish Neo UHPLC system. The instrument operated at a flow rate of 0.4 *µ*l min-1 with buffer A (97.9% Water, 2% Acetonitrile, and 0.1% Formic Acid) and buffer B (97.9% Acetonitrile, 2% Water, and 0.1% Formic Acid). Peptides were separated by a multi-step gradient: 0-1 min from 2% B to 4% B, 1-44 min from 4% B to 22.5% B, 44-59 min from 22.5% B to 35% B, followed by a column wash. One microliter of each K562 and CSF dilutions were injected for the LC–MS/MS analysis.

The Orbitrap Astral Mass Spectrometer was operated in a positive scan DIA mode at a full-MS resolution of 240,000 with a scan range of 380–980 m/z. The normalized AGC target was set to 500%. The MS/MS scans were recorded in a profile mode at a maximum injection time of 5 ms, 4Th scanning from 380 to 980 m/z. HCD with 25% normalized collision energy was used to fragment the ions.

##### Spectronaut data analysis

Raw files from K562 and CSF dilutions benchmark dataset were analyzed in Spectronaut v19 (Biognosys) with a library-free approach (directDIA+). The human reference database (UniProt, 20,594 sequences) was utilized for this analysis. For modifications, cysteine carbamydomethylation was set as a fixed modification. Methionine oxidation and protein N-terminal acetylation were set as variable modifications. Precursor filtering was set to a 0.01 Q value, and cross-run normalization was unchecked.

##### Statistical analysis

In both experiments, two conditions were generated by mixing the same material in two different ratios to simulate a 2-fold change in protein abundance across all proteins. To simulate a realistic discovery proteomics experiment, we randomly selected 90% of the quantified proteins and swapped their condition labels, eliminating any difference between conditions for those proteins. Within each condition, 15 samples used consistent sample input, representing ideal conditions. The remaining 5 samples in each group were intentionally diluted at progressively lower concentrations to simulate instrumental drift. The specific mixture ratios, run annotations, and swapped condition labels are provided in **Supplementary Table 7** and **Supplementary Table 8**.

##### Mixture of proteomes

Mixed-species proteome samples containing human, yeast, and Escherichia coli proteins in six different ratios were analyzed using an Orbitrap Astral Mass Spectrometer operating in DIA mode. Detailed information about the experimental setup and data acquisition can be found in Guzman et al. [49]. The resulting datasets were downloaded from ProteomeXchange (identifier PXD046444) in April 2025 and processed using Spectronaut v19. This experiment, which quantified over 14,000 protein groups, was specifically designed by the authors to demonstrate the superior label-free quantification accuracy and precision of their optimized workflow for the Astral analyzer.

This workflow combines a narrow-window data-independent acquisition (nDIA) strategy with high-resolution MS1 scans and parallel ultra-fast MS2 scans at 200 Hz, utilizing 2Th isolation windows. By minimizing the co-elution of multiple peptides and reducing chimeric spectra (a common issue in traditional DIA methods), this approach is particularly well-suited for applications that traditionally rely on DDA strategies. Additionally, it still addresses the missing value problem commonly associated with conventional DDA workflows. As a result, it achieves threefold higher proteome coverage in a fraction of the time required by current state-of-the-art mass spectrometry methods, offering a robust workflow for highthroughput, comprehensive proteome profiling.

Unlike the benchmark K562 and CSF datasets, this experiment inherently featured low noise and very few missing values, with no intentionally introduced anomalous measurements.

##### Clinical cohort

This experiment reflected the complexity of real-world clinical proteomics, including large number of samples, biological variability, instrumental drift, and missing values. The study investigated the safety and efficacy of Semorinemab in Alzheimer’s disease. CSF samples were collected from patients in two clinical trials studying Alzheimer’s disease: Tauriel (at baseline and after 49 or 73 weeks) and Lauriet (at baseline and after 49 or 61 weeks). Samples were analyzed using single-shot FAIMS-DIA-MS on an Orbitrap Exploris 480 Mass Spectrometer and the data was acquired using Spectronaut v19. Proteins were quantified in a broad dynamic range with intensities spanning 7 orders of magnitude. Detailed information about the experimental setup and data acquisition can be found in Abdel-Haleem et al. [51]

### 4.2. MSstats+ workflow

#### Step 1: Extract fragment peak intensities and precursor quality metrics

*MSstats+* supports a variety of experimental designs (e.g., group comparison, time course, paired designs) and unbalanced designs (i.e., with unequal number of replicates in each condition). *MSstats+* takes as input (1) fragment-level intensity measurements, (2) precursor-level spectral quality metrics, (3) longitudinal order of MS run acquisition, (4) the peptide and protein identifiers of the precursor ions reported by a data processing tool, and (5) metadata describing conditions, subjects and other annotations of each run.

For a given protein, denote 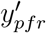 the intensity of precursor *p* = 1, …, *P*, fragment *f* = 1, …, *F*, and run *r* = 1, …, *R*. Similarly, denote 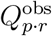 a vector of all the observed spectral quality metrics of precursor *p* and run *r*. We use “*·*” in place of the fragment subscript to indicate that the metric aggregates the measurements over the fragments. The metrics can be reported by any upstream data processing tool. We recommend using metrics known to be informative of the quality of the quantification. Alternatively, it is possible to explore each metric empirically and retain the metrics where Pearson’s moment coefficient of skewness shifts away from 0 in at least some precursors.

#### Step 2: Filter, transform and normalize fragment intensities

Step 2 performs univariate feature filtering designed to remove fragments and peak intensities that are obviously low quality, and ensures that the remaining low quality measurements are relatively rare. These options depend on the information provided by the data processing tool and can vary based on if the rows are removed entirely (including the quality metrics), or if the intensity values are set to NA (retaining the quality metrics). The exact filtering steps for Spectronaut are listed below.

First, remove measurements with any type of fragment loss. Fragment losses are anno-tated across the entire fragment for all runs, and these rows are removed from the dataset. Second, set intensity values to `NA` if Spectronaut marks them as excluded from quantification. Spectronaut flags such values in the column `F.ExcludedFromQuantification.` Any fragment-level intensity measurement with `F.ExcludedFromQuantification = TRUE` is set to `NA.`

Third, set intensity values to `NA` if they do not pass the q-value filtering threshold. Qvalues are reported at both the protein group level (`PG.Qvalue`) and the elution group level (`EG.Qvalue`). Any measurement with a q-value greater than 0.01 at either level is set to `NA`. Fourth, set measurement intensities below 1 to `NA`. Spectronaut rarely reports missing values (`NA`) in the `F.PeakArea` or `F.PeakHeight` columns. However, values below an intensity of 1 represent background noise rather than true signal and are treated as missing.

Fifth, balance the experimental design by adding additional rows with `NA` intensities and quality metrics. For each fragment, we check whether it is represented in all runs. If a fragment is missing in any run, a new row is added for that run with `NA` values for both intensity and quality metrics.

Sixth, filter out missing fragments in more than 50% of runs. After the previous steps, many intensities may have been set to `NA`. Here, we remove fragments observed in less than half of the runs (threshold adjustable by the user).

Seventh, optionally perform top-N feature filtering using a large number of the highest-intensity fragments (e.g., top-100). In large DIA experiments, some proteins are represented by thousands of fragments, which slows computation without adding meaningful information (as seen in the comparison between top-100 and all features in **Supplementary Fig. 1**) [50]. This step provides broad intensity-based feature filtering while retaining a large number of top intensity fragments.

Eighth, apply quantile filtering to remove intensities near the LoD or LoQ. Intensities near these thresholds are generally less reliable. We therefore set to `NA` all intensities falling within a small lower quantile (default: below the 0.1 percentile) of all measurements [30].

Finally, we apply standard data processing steps, including log transformation of intensities such that 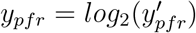, and normalization using the methods described in Kohler et al [30].

#### Step 3: Engineer longitudinal transformations of quality metrics; model quality metrics into a single anomaly score

##### Engineer longitudinal transformations of quality metrics

To capture drifts in peak quality over time, *MSstats+* engineers additional longitudinal quality metrics of each precursor [43], denoted 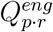. The features are based on three CUSUM statistics [55, 67]. The constants in the statistics are derived from the assumption that the quality metrics are centered and scaled separately for each precursor, and that they are Normally distributed when measurements are collected under optimal performance.

Positive CUSUM (mean increase) captures increasing trends in mean quality. Denote *q*_*r*_ the value of one dimension of the observed vector of quality metric of a precursor in run *r* (for simplicity of notation, we omit the remaining subscripts). Define *S*_1_ = 0. Then for *r ≥* 2,

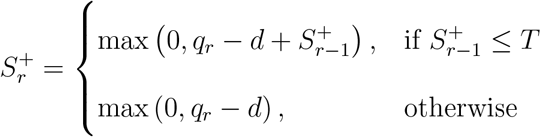

where *d* = 0.5 is the allowable drift and *T* = 5 is the reset threshold. Negative CUSUM (mean decrease) captures decreasing trends in mean quality. Define *S*_1_ = 0. Then for *r ≥* 2,

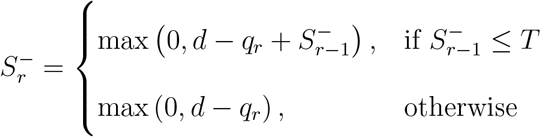

with *d* = *−*0.5 and the same reset threshold *T* = 5. CUSUM for dispersion increase captures increase in variability. We first introduce an additional normalization of the square root of each quality value

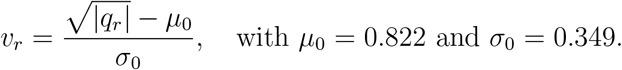

[68]. Then, using the same recurrence form as above, set *D*_1_ = 0 and define

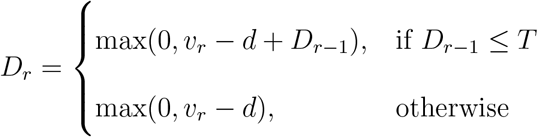

with *d* = 0.5. We recommend applying only one transformation per metric. For example, if we have prior knowledge that a metric will decrease over time, only the negative CUSUM should be used (e.g., peak quality score). Conversely, if we anticipate random shifts in both directions, then CUSUM for dispersion increase should suffice (e.g., delta retention time). When no prior knowledge is available, we recommend defaulting to using CUSUM for dispersion increase, as it can detect both upward and downward changes.

After calculating engineered CUSUM quality metrics for all precursors P and runs R, the metrics are concatenated into a full vector 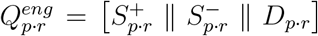 . Finally, *MSstats+* concatenates the original and the engineered metrics into a full vector 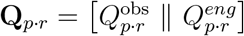 to capture both static and longitudinal characteristics of peak quality for the downstream analysis.

##### Model quality metrics into a single anomaly score

Next, *MSstats+* transforms the vector of observed and engineered metrics **Q**_*p·r*_ into a single anomaly score with isolation forest (iForest). *MSstats+* uses a custom implementation of iForest specifically designed for MS-based proteomics data. A separate iForest is built for each precursor *p*. It views each run as a multivariate observation of quality metrics for the precursor, and constructs *N* random binary trees *T*_*p*;*n*_ (*T*_*p*;*n*_ = 100 as a default). Each tree is generated by iteratively randomly sampling one quality metric dimension and one split value in that dimension. Runs are split into two groups based on the sampled split value. The implementation incorporates the presence of missing anomaly metrics (unreported by Spectronaut, or introduced by the unbalanced nature of the experimental design) as an additional feature for tree splitting. This process is repeated until the tree reaches its maximum depth (default [*log*_2_(*R*)]). After building the forest, the average path length to each run is calculated as:

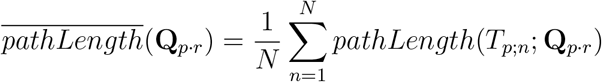

The anomaly score *a*_*p·r*_ is then computed following [48] as:

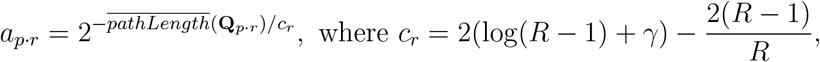

*R* is the number of runs and *γ* is Euler’s constant. Anomaly scores *a*_*p·r*_ are bounded between 0 and 1 and can be viewed as a probability. Anomaly scores near 0 correspond to high-quality measurements, while scores near 1 indicate poor quality.

#### Step 4: Summarize weighted fragment intensities into protein-level abundance

A separate protein-level summarization is performed for each protein. It takes as input all fragments from all precursors that remain in the dataset after univariate filtering and pertain to the protein. Missing fragment intensities are imputed as implemented in *MSstats*, using the Accelerated Failure Time (AFT) model [69] described in Kohler *et al* [30]. Anomaly scores *a*_*p·r*_ are calculated for both observed and imputed fragment intensities. See **Supplementary Section 2.10** for examples of specific scenarios.

Similarly to *MSstats*, each protein summarizes the log_2_ intensities *y*_*pfr*_ of precursors *p* = 1, …, *P*, fragments *f* = 1, …, *F* and runs *r* = 1, …, *R* in a regression model

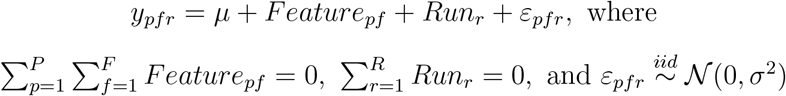

Unlike *MSstats*, parameter estimation in *MSstats+* weights each fragment intensity by the inverse of its precursor-level anomaly score. In practice, we form weights from anomaly scores by lower-bounding the score before inversion, such that

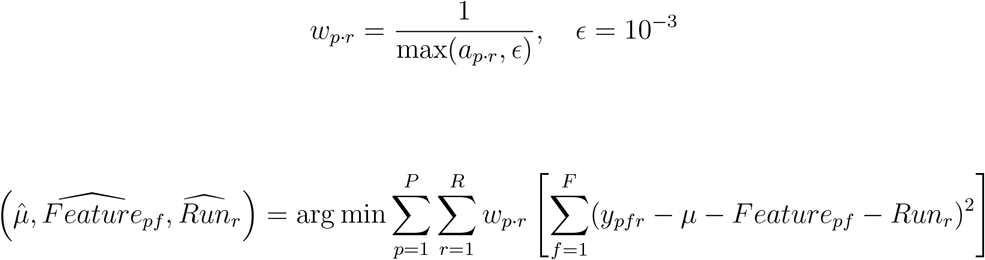

Finally, protein-level summary of run *r* is 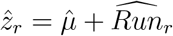. Its variance 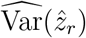 is derived according to the standard procedure for weighted regression [70]. The variance serves as a measure of uncertainty associated with the summary in that run.

#### Step 5: Model weighted protein-level abundance and perform differential analysis

Similarly to *MSstats, MSstats+* fits a separate linear mixed effects model to each protein summary 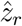. The model includes terms that describe the systematic variation in protein-level summaries induced by the experimental design. *MSstats+* automatically selects the appropriate terms for each design. The model also includes terms that reflect the nonsystematic variation between runs. Unlike in *MSstats*, these terms account for the unequal uncertainty associated with 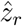. For example, in a group comparison design with no technical replicates, *MSstats+* fits the model

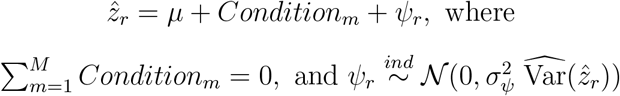

The terms *µ* and *Condition*_*m*_ reflect the systematic variation, and the term *ψ*_*r*_ reflects the non-systematic variation between runs. The index *m* = 1, …, *M* maps each run *r* to one of the *M* conditions. Similarly to Step 4, parameters of the model are estimated by weighted least squares (or weighted restricted maximum likelihood for more complex designs [71]). For example, for the model above the estimation is

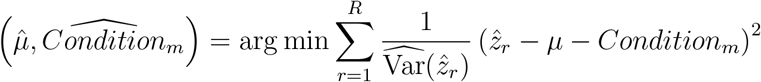

The variance 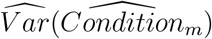 is derived according to the standard procedure for weighted regression [70], and reflects the uncertainty associated with the estimation. Model-based testing of hypotheses of interest is then performed using contrasts as described in Kohler *et al*. [30]. For example, in the model above, a comparison between Condition 2 versus Condition 1 proceeds by comparing

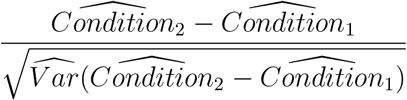

to the Student distribution with the model-derived degrees of freedom.

## Supporting information

Supplemental manuscript

## 5. Data Availability

The raw data for the three-species mix benchmark study can be found on ProteomeXchange under the identifier PXD046444 [49]. The Spectronaut file for the three-species mix benchmark study, along with the K562 and CSF MS and search data, has been deposited in MassIVE under the identifier MSV000098622 and can also be accessed through ProteomeXchange with the identifier PXD066486. The statistical results for the clinical study can be found in **Supplementary Data 1**.

## 6. Code Availability

The implementation and instructional vignette for *MSstats+* are available on Github at github.com/Vitek-Lab/MSstats and github.com/Vitek-Lab/MSstatsConvert, with planned release on Bioconductor (October 2025). The analysis code and intermediate data files to reproduce the results in this manuscript are available in MassIVE under RMSV000000701.2.

