## Supplemental manuscript for "Accounting for longitudinal peak quality metrics with MSstats+ enhances differential analysis in proteomic experiments with data-independent acquisition"

### Supplementary Information

Devon Kohler[1], Eralp Dogu[2], Mrittika Bhattacharya[3], Ozge Karayel[3], Manuel Magana[3], Anthony Wu[1], Veronica Anania[3], Olga Vitek[1*]

[1] Khoury College of Computer Sciences and Barnett Institute for Chemical and Biological Analysis, Northeastern University, Boston, MA, USA 02115
[2] Mugla Sitki Kocman University, Department of Statistics, Mugla, Turkiye
[3] Genentech, South San Francisco, CA, USA 94080

### 1 Introduction

#### 1.1 Using a low number of features in Top-N feature selection increases the FDR in a benchmark experiment

Standard feature selection algorithms use intensity (or intensity derived metrics such as feature variation) to determine which features best represent the underlying protein. While these algorithms can be effective at reducing variation in the experiment they can also introduce a double dipping problem, where intensities are used to select features and then those same intensities are used in differential analysis. Here we test top-N feature selection at 6 different levels and record the false discovery rate (FDR) in a DIA benchmark experiment measured by Navarro et al [1].

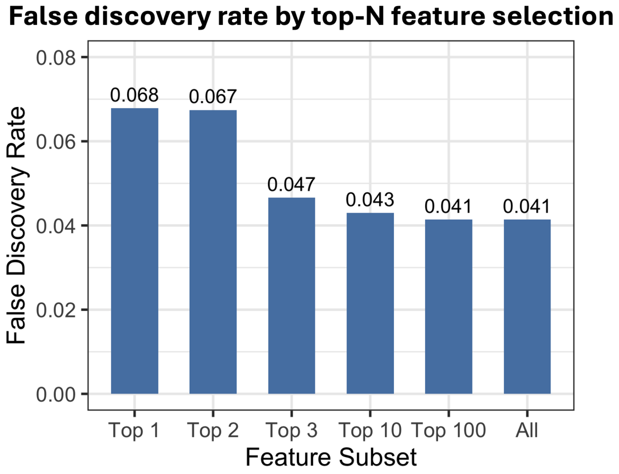

Supplementary Fig. 1: **Lower top-N feature selection is associated with higher FDR in a DIA benchmark experiment by Navarro et al [1].** The y-axis shows the false discovery rate (FDR), and the x-axis indicates the number of top features selected per protein. The experiment was processed using standard *MSstats* with a nominal FDR cutoff of 0.05, while varying numbers of top most abundant features used for protein-level summarization. Using only the top 1 or 2 features resulted in the highest FDR, exceeding 6%. Selecting the top 3 features reduced FDR, but it remained higher than the nominal level. FDR stabilized once Top-N exceeded 100, with no difference between top 100 features and all features.

#### 1.2 Individual filtering cutoffs are unable to separate high quality and low quality measurements in an experiment with known run labels

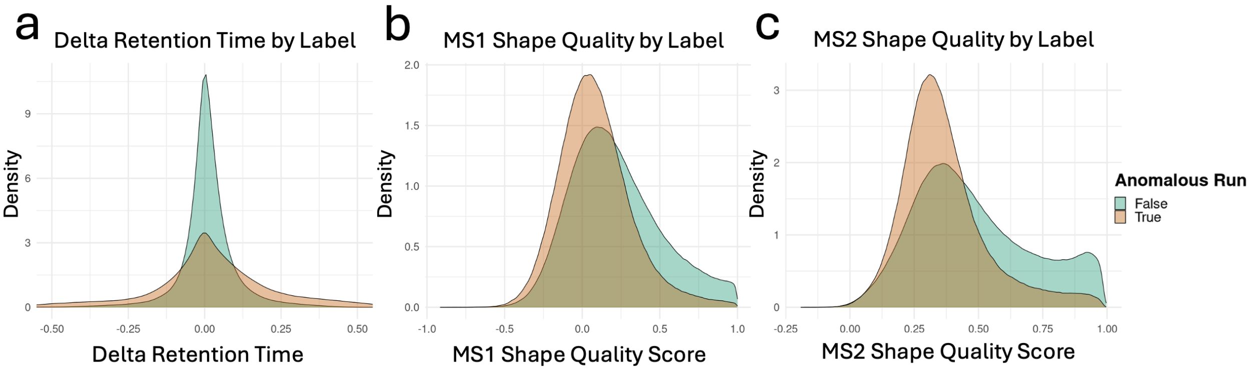

Supplementary Fig. 2: **Dataset 1: K562 benchmark. Distribution of spectral peak quality metrics cannot separate high quality and low quality measurements with one filtering value.** Distributions of delta retention time (a), MS1 shape quality score (b), and MS2 shape quality score (c), subset by known anomalous (orange) and non-anomalous (green) runs. While there are clear shifts in the distributions between anomalous and non-anomalous runs, no quality metric provides a clean separation, making it difficult to define a universal filtering threshold. Beyond this, any single filtering threshold will inevitably include poorly quantified measurements in the final output, while also removing high quality measurements.

### 2 Results

#### 2.1 Example Spectronaut output file shows where standard filtering steps are required

| | R.Run Date (Formatted) | R.Empirical MS1 Resolution | R.Condition | R.FileName | R.Replicate | PG.ProteinAccessions | PG.ProteinGroups | PG.Qvalue | PG.Quantity | PEP.GroupingKey | PEP.StrippedSequence | PEP.Quantity |
|---|---|---|---|---|---|---|---|---|---|---|---|---|
| | <char> | <char> | <char> | <char> | <int> | <char> | <char> | <num> | <num> | <char> | <char> | <num> |
| 1: | 1/3/2025 9:03:40 PM | 223,400 | 1to2 | 20250102_Tulum_1to2K562_Seq7 | 3 | P48059 | P48059 | 0 | 83.962051 | GGFAPAEK | GGFAPAEK | 305.975952 |
| 2: | 1/4/2025 7:51:32 AM | 223,900 | Neat | 20250102_Tulum_NeatK562_Seq15 | 8 | Q6IN85 | Q6IN85 | 0 | 105.429214 | AFLNNGSR | AFLNNGSR | 140.213486 |
| 3: | 1/5/2025 4:17:25 PM | 227,100 | 1to64 | 20250102_Tulum_1to64K562_Seq39 | 1 | Q8N108 | Q8N108 | 0 | NaN | DNEQALYELVK | DNEQALYELVK | NaN |
| 4: | 1/4/2025 1:15:42 PM | 224,000 | 1to2 | 20250102_Tulum_1to2K562_Seq19 | 9 | P53602 | P53602 | 0 | 168.254745 | IIHLVHR | IIHLVHR | 1.987759 |
| 5: | 1/5/2025 8:10:52 AM | 223,900 | 1to8 | 20250102_Tulum_1to8K562_Seq33 | 1 | O95671 | O95671 | 0 | 8.055086 | VLCPVIGK | VLCPVIGK | NaN |

| | EG.iRTPredicted | EG.Library | EG.ModifiedSequence | EG.PrecursorId | EG.Qvalue | EG.ApexRT | EG.DeltaRT | EG.FWHM | EG.PeakWidth | FG.ShapeQualityScore (MS1) | FG.ShapeQualityScore (MS2) |
|---|---|---|---|---|---|---|---|---|---|---|---|
| | <num> | <char> | <char> | <char> | <num> | <num> | <num> | <num> | <num> | <num> | <num> |
| 1: | -21.68894 | directDIA™ | _GGFAPAEK_ | _GGFAPAEK_.2 | 7.880308e-11 | 11.24834 | 0.015995177 | 0.07610033 | 0.10407543 | 0.2351272 | 0.7702653 |
| 2: | -12.22753 | directDIA™ | _AFLNNGSR_ | _AFLNNGSR_.2 | 1.606263e-08 | 14.73504 | -0.001698438 | 0.04658490 | 0.09096909 | 0.3551088 | 0.4165565 |
| 3: | 54.28896 | directDIA™ | _DNEQALYELVK_ | _DNEQALYELVK_.2 | 7.554978e-02 | 40.04576 | -0.088275203 | 0.02021444 | 0.03643417 | 0.2004314 | 0.2271121 |
| 4: | -13.61469 | directDIA™ | _IIHLVHR_ | _IIHLVHR_.2 | 2.483772e-03 | 14.35648 | 0.288184307 | 0.12448474 | 0.03651333 | 0.2399451 | 0.2672812 |
| 5: | 16.98906 | directDIA™ | _VLC[Carbamidomethyl (C)]PVIGK_ | _VLC[Carbamidomethyl (C)]PVIGK_.2 | 2.535252e-02 | 24.93583 | -0.334040633 | 0.10761820 | 0.07895851 | 0.1939022 | 0.2345202 |

| | FG.MS1IsotopeIntensities (Measured) | FG.MS1IsotopeRelativeIntensities (Measured) | F.PriorIonRatio | FG.Charge | FG.Id | FG.PrecMz | FG.ShapeQualityScore | FG.MS2RawQuantity |
|---|---|---|---|---|---|---|---|---|
| | <char> | <char> | <num> | <int> | <char> | <num> | <num> | <num> |
| 1: | 94523.6;26653.6;25703.6;18484.5 | 0.572;0.161;0.155;0.112 | 0.001826666 | 2 | _GGFAPAEK_.2 | 388.7005 | 0.6200429 | 305.975952 |
| 2: | 69748.8;45181.9;31904.6;24437.8 | 0.407;0.264;0.186;0.143 | 0.132126153 | 2 | _AFLNNGSR_.2 | 439.7276 | 0.3779793 | 140.213486 |
| 3: | 0.8;0.5;0.9;0.9;0.9 | 0.204;0.126;0.227;0.214;0.229 | 0.100728683 | 2 | _DNEQALYELVK_.2 | 661.3353 | 0.1784342 | 5.195623 |
| 4: | 26230.4;1;0.9;0.9 | 1;0;0;0 | 0.024863454 | 2 | _IIHLVHR_.2 | 444.2823 | 0.1961655 | 1.987759 |
| 5: | 68557.4;22170.4;6676.2;0.8 | 0.704;0.228;0.069;0 | 0.071360454 | 2 | _VLC[Carbamidomethyl (C)]PVIGK_.2 | 443.2650 | 0.2078423 | NaN |

| | FG.HasPossibleInterference (MS1) | FG.HasPossibleInterference (MS2) | FG.Quantity | FG.CalibratedMassAccuracy (PPM) | FG.Noise | FG.RawMassAccuracy (PPM) | F.Charge | F.FrgIon | F.FrgLossType | F.FrgMz | F.FrgNum | F.FrgType |
|---|---|---|---|---|---|---|---|---|---|---|---|---|
| | <lgcl> | <lgcl> | <num> | <num> | <num> | <num> | <int> | <char> | <char> | <num> | <int> | <char> |
| 1: | TRUE | TRUE | 305.975952 | 0.06427352 | 751.7352 | -1.7272598 | 1 | y4 | noloss | 444.2453 | 4 | y |
| 2: | TRUE | TRUE | 140.213486 | -1.19081960 | 374.1998 | -1.5268242 | 1 | y5 | NH3 | 530.2318 | 5 | y |
| 3: | TRUE | TRUE | 5.195623 | NaN | 420.3224 | NaN | 1 | y7 | noloss | 835.4924 | 7 | y |
| 4: | FALSE | TRUE | 1.987759 | 1.57323385 | 368.4425 | 0.1373792 | 2 | y5 | noloss | 331.1983 | 5 | y |
| 5: | TRUE | TRUE | NaN | -0.96608014 | 194.7357 | -1.1704035 | 1 | b3 | noloss | 373.1904 | 3 | b |

| | F.ExcludedFromQuantification | F.NormalizedPeakArea | F.NormalizedPeakHeight | F.PeakArea | F.PeakHeight | F.CalibratedMassAccuracy (PPM) | F.CalibratedMz | F.HasChannelInterference | F.InterferenceScore |
|---|---|---|---|---|---|---|---|---|---|
| | <lgcl> | <num> | <num> | <num> | <num> | <num> | <num> | <lgcl> | <num> |
| 1: | FALSE | NaN | NaN | 144.66387939 | 2950.1401367 | 0.9578953 | 444.2440 | FALSE | 0.00000000 |
| 2: | FALSE | NaN | NaN | 0.09096909 | 0.9965138 | NaN | 530.2304 | FALSE | 0.00000000 |
| 3: | FALSE | NaN | NaN | 0.03643417 | 0.6715688 | NaN | 835.4908 | FALSE | 0.25519025 |
| 4: | FALSE | NaN | NaN | 1.91473246 | 97.1105804 | -2.0174803 | 331.1973 | FALSE | 0.01039139 |
| 5: | TRUE | NaN | NaN | 0.66089463 | 64.8150558 | NaN | 373.1897 | FALSE | 0.39113429 |

| | F.IsotopicPatternTheoretical | F.Log10SignalToNoise | F.MSPeakApexMeasuredMz | F.Noise | F.PossibleInterference | F.PPMTolerance | F.MassAccuracyPPM | F.TheoreticalMz | F.Tolerance |
|---|---|---|---|---|---|---|---|---|---|
| | <char> | <num> | <num> | <num> | <lgcl> | <num> | <num> | <num> | <num> |
| 1: | 1;0.244;0.031 | NaN | NaN | NaN | FALSE | 4.408582 | 3.7782435 | 444.2453 | 0.001958492 |
| 2: | 1;0.285;0.049 | NaN | NaN | NaN | FALSE | 3.492764 | NaN | 530.2318 | 0.001851975 |
| 3: | 1;0.434;0.124;0.027 | NaN | NaN | NaN | FALSE | 8.591916 | NaN | 835.4924 | 0.007178480 |
| 4: | 1;0.348;0.079;0.015 | NaN | NaN | NaN | FALSE | 5.160800 | 0.8292863 | 331.1983 | 0.001709248 |
| 5: | 1;0.211;0.018 | NaN | NaN | NaN | TRUE | 4.914968 | NaN | 373.1904 | 0.001834219 |

Supplementary Fig. 3: **Example 5 rows from the Spectronaut PSM output file for Dataset 1: K562 benchmark, highlighting key filtering steps.** In step 1 of filtering, row 2 will be removed from the data due to the "NH3" value in `F.FrgLossType`. In step 2 of filtering, Row 5 has a TRUE value in `F.ExcludedFromQuantification`, indicating the intensity for this row will be set to `NA`. In step 3, all of the values in both `PG.Qvalue` and `EG.Qvalue` are below 0.01, meaning these rows will not be filtered out. In step 4, Rows 2 & 3 have a `F.PeakArea` below 1, indicating they will be treated as missing values in the filtering steps and set to `NA`. Beyond filtering columns, we can see example quality metrics which can be used in the anomaly detection model: `FG.ShapeQualityScore (MS1)`, `FG.ShapeQualityScore (MS2)`, `EG.ApexRT`, `EG.DeltaRT`, and `EG.PeakWidth`.

#### 2.2 Fragment-level anomaly model is less sensitive and increases computational runtime in benchmark studies

Spectronaut reports spectral peak quality metrics on the elution group, precursor group, and individual fragment levels. We contrast two anomaly models: one using only PSM-level quality metrics, and another that incorporates both PSM and fragment-level quality metrics. We compare the performance of each model in terms of their ability to correctly identify true positives ($\alpha = 0.05$) and their computational runtime. The models are evaluated on both Dataset 1: K562 benchmark and Dataset 2: CSF benchmark.

The PSM-level model included the following quality metrics: MS1 peak quality, MS2 peak quality, delta retention time. The fragment-level model used the same metrics as the PSM-level model and additionally included PPM tolerance, and calibrated mass accuracy.

| | Dataset 1: K562 benchmark | | Dataset 2: CSF benchmark | |
|---|---|---|---|---|
| | PSM-level | Fragment-level | PSM-level | Fragment-level |
| TPR | 0.914 | 0.768 | 0.825 | 0.006 |
| Runtime (minutes) | 24.33 | 47.66 | 6.010 | 10.933 |

Supplementary Table 1: **Performance comparison of PSM and fragment-level anomaly models.** True positive rate (TPR, $\alpha = 0.05$) and runtime are reported for both PSM-level and fragment-level models. All benchmarks were parallelized across 24 cores using an Intel(R) Xeon(R) CPU E5-2680 v4 @ 2.40GHz with 24 compute nodes. Across both benchmarks, the PSM-level model outperformed the fragment-level model in both TPR and runtime.

#### 2.3 C++ implementation of the the isolation forest improved computational processing time

a

| Model | 1 Core | 12 Cores |
|---|---|---|
| R | 1.66h | 7.63m |
| C++ | 4.48m | 45.07s |

b

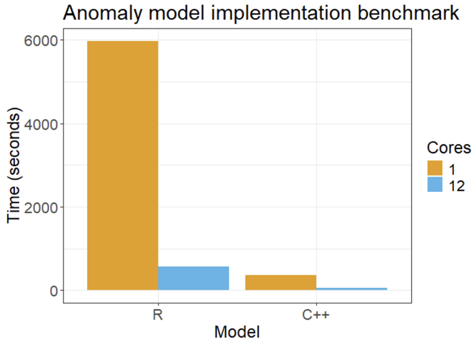

Supplementary Fig. 4: **Dataset 1: K562 benchmark (1000 protein subset). C++ implementation with parallel processing greatly improved computational processing time of isolation forest model.** All benchmarks were run on an Intel(R) Xeon(R) CPU E5-2680 v4 @ 2.40GHz. Two views of the computational time with **(a)** exact values in a table and **(b)** a bar plot showing the scale of differences. Using R with one core was the slowest implementation. Adding parallel processing to the R implementation and using 12 CPU cores reduced the computational time by 92.3%. Implementing the model in C++ improved the time over both R implementations ( 41.2% decrease over the 12 core R implementation). The parallelized C++ implementation with 12 cores further reduced the processing time by 83.2% over the single core C++ version. Overall, using both C++ and parallel processing greatly reduced the computational time.

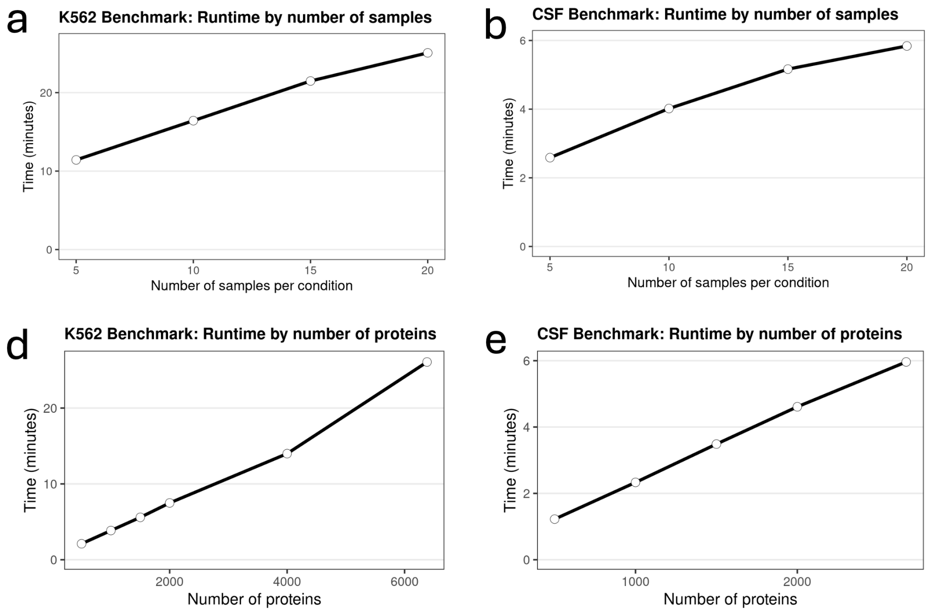

Supplementary Fig. 5: **K562 and CSF benchmarks *MSstats+* runtime scales when increasing the number of proteins and runs.** All benchmarks were performed using an Intel(R) Xeon(R) CPU E5-2680 v4 @ 2.40GHz with 24 compute nodes. **(a)** K562 benchmark runtime while varying the number of samples per condition using all proteins. The runtime remains relatively high even with few samples and scaled approximately linearly as samples increased. **(b)** Same as (a) for the CSF benchmark. **(c)** K562 benchmark runtime while varying the number of proteins per condition using all samples. The runtime was much lower with a small number of proteins compared to a low numbers of samples. As the number of proteins increased, the runtime quickly increased. This is caused by the need to fit a new iForest for each additional precursor. **(d)** Same as (c) for the CSF benchmark.

#### 2.4 Additional peak quality metrics for A4D126 improve differential analysis results compared to traditional methods

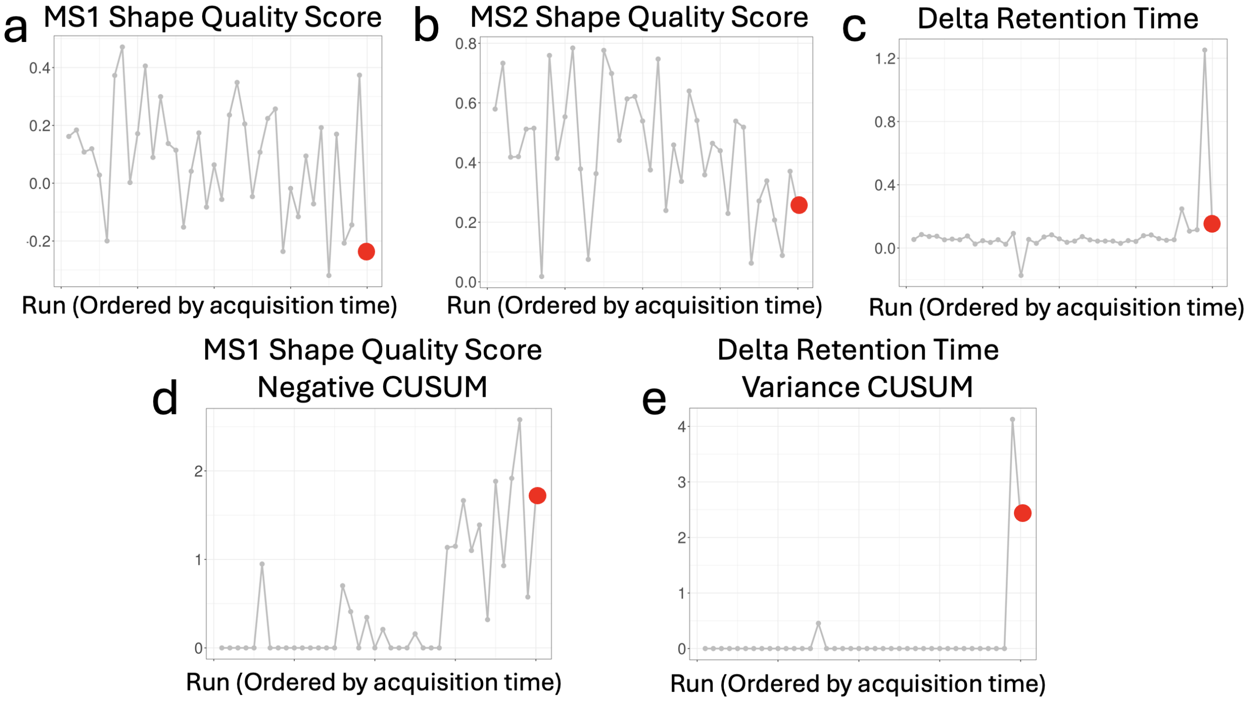

Supplementary Fig. 6: **Dataset 1: K562 benchmark. Precursor-level quality metrics for protein A4D126 and precursor ALAEDQINSK.2 show a drop in quality in run 40.** (**a**) The MS1 shape quality scores across all runs. Run 40 is marked in red. The MS1 shape quality score sees a substantial drop in run 40 compared to the average across all runs. (**b**) Same as (a) for MS2 shape quality score. Again the MS2 shape quality score is lower in run 40 compared to the average across all runs. (**c**) Same as (a) for delta retention time. The retention time across all runs is mostly stable, with one big increase in run 39. Run 40 appears mostly in line with the majority of runs. (**d**) Same as (a) for the engineered MS1 shape quality negative CUSUM metric. This metric captures downward trends in shape quality score at the MS1 level. Near the end of the run order, the metric shows a sharp increase which remains high until run 40. This captures a drop in quality trend as runs were acquired at the MS1 level. (**d**) Same as (a) for the engineered delta retention time variance CUSUM metric and captures variation in the delta retention time. This metric is zero in nearly every run but shows a large increase in run 39 due to the jump in delta retention time seen in (c). This increase influenced run 40 which also remained higher than the other runs, even though it was low in (c).

| **Model** | **$\text{Log}_2$ Fold Change** | **Standard Error** | **Adj P-value** |
|---|---|---|---|
| MSstats+ | -1.110 | 0.321 | 0.017 |
| MSstats | -0.769 | 0.502 | 0.999 |
| msqrob2 | -1.033 | 0.592 | 0.936 |
| limma | -0.824 | 0.345 | 0.525 |
| limpa | -0.951 | 0.506 | 0.920 |
| DEqMS | -0.745 | 0.324 | 0.674 |

Supplementary Table 2: **Differential analysis for a single protein (A4D126) in Dataset 1: K562 benchmark across all statistical methods evaluated.** The true $\log_2$ fold change is -1. The proposed approach was the only method to achieve a differentially abundant adjusted p-value. Across all methods, the proposed approach returned the lowest standard error due to down-weighting anomalous runs.

#### 2.5 Additional protein examples show effectiveness of MSstats+ in the presence of known anomalous runs

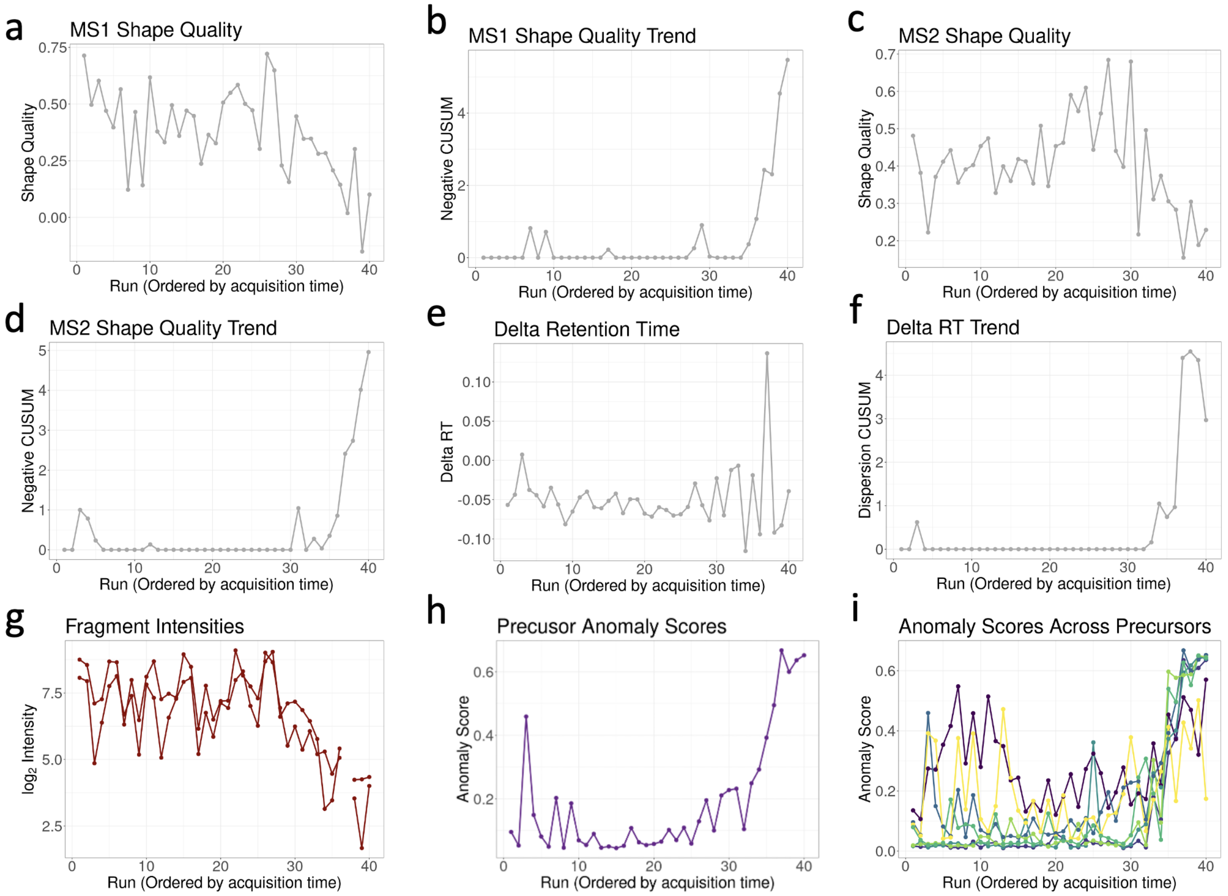

Supplementary Fig. 7: **Dataset 1: K562 benchmark. Precursor-level quality metrics and intensities for protein Q9Y5S9 and precursor EDYDSVEQDGDEPGPQR.2.** *MSstats+* detected the manually introduced anomalous samples after run 30, as well as a singular jump in anomaly score in run 3. (**a**) MS1 shape quality scores across runs. Scores are generally high before run 30, with smaller drops in runs 7 and 9. (**b**) MS1 shape quality negative CUSUM. (**c**) MS2 shape quality score. One clear drop occurs in run 3. (**d**) MS2 shape quality negative CUSUM. (**e**) Delta retention time. The metric is generally stable, without any outliers, until after run 30, when the variance of the measurements increases. (**f**) Delta retention time dispersion CUSUM. The increase in variance seen in (e) is observed here. (**g**) Precursor intensities, with each line representing a different fragment. (**h**) The anomaly score in each run. Run 3 shows a clear increase, caused by the drop in MS2 shape quality in (c). There are also small increases in runs 7 and 9 due to drops in MS1 quality in (a). (**i**) The anomaly score across all precursors for protein Q9Y5S9. All precursors show an increase in anomaly scores after run 30.

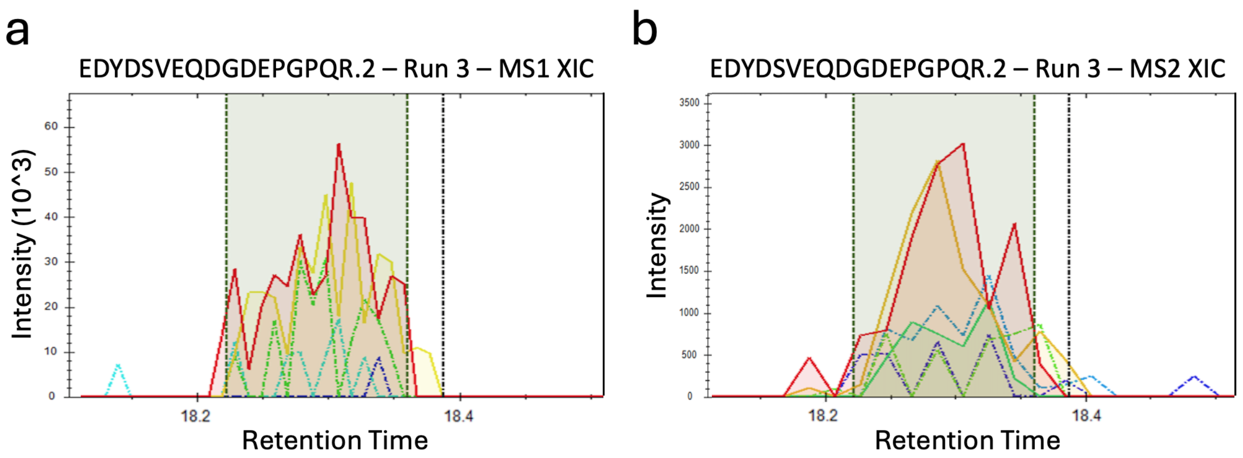

Supplementary Fig. 8: **Dataset 1: K562 benchmark. MS1 and MS2 extracted ion chromatogram (XIC) visualizations for protein Q9Y5S9 and precursor EDYDSVEQDGDEPGPQR.2 in Run 3.** In **Supplementary Fig. 7**, *MSstats+* detected a singular jump in anomaly score in run 3, which can be manually validated using the XIC. The predicted retention time is indicated by the black dotted line, and the observed retention window is highlighted in green. (**a**) MS1 XIC for EDYDSVEQDGDEPGPQR.2. Each colored line represents an isotopic peak of the precursor. The MS1 XIC peak shape appeared to be high quality with the isotopic peaks broadly aligned. This corresponds to the observation in **Supplementary Fig. 7**a, where the MS1 shape quality was high in run 3. (**b**) MS2 XIC for EDYDSVEQDGDEPGPQR.2. Each colored line represents a fragment ion of the precursor. The MS2 XIC showed clearly defined fragment peaks, however they were not perfectly aligned and the y5 ion (colored red) was not Gaussian and contained two peaks. This corresponds to the observation in **Supplementary Fig. 7**c, where the MS2 shape quality was low in run 3.

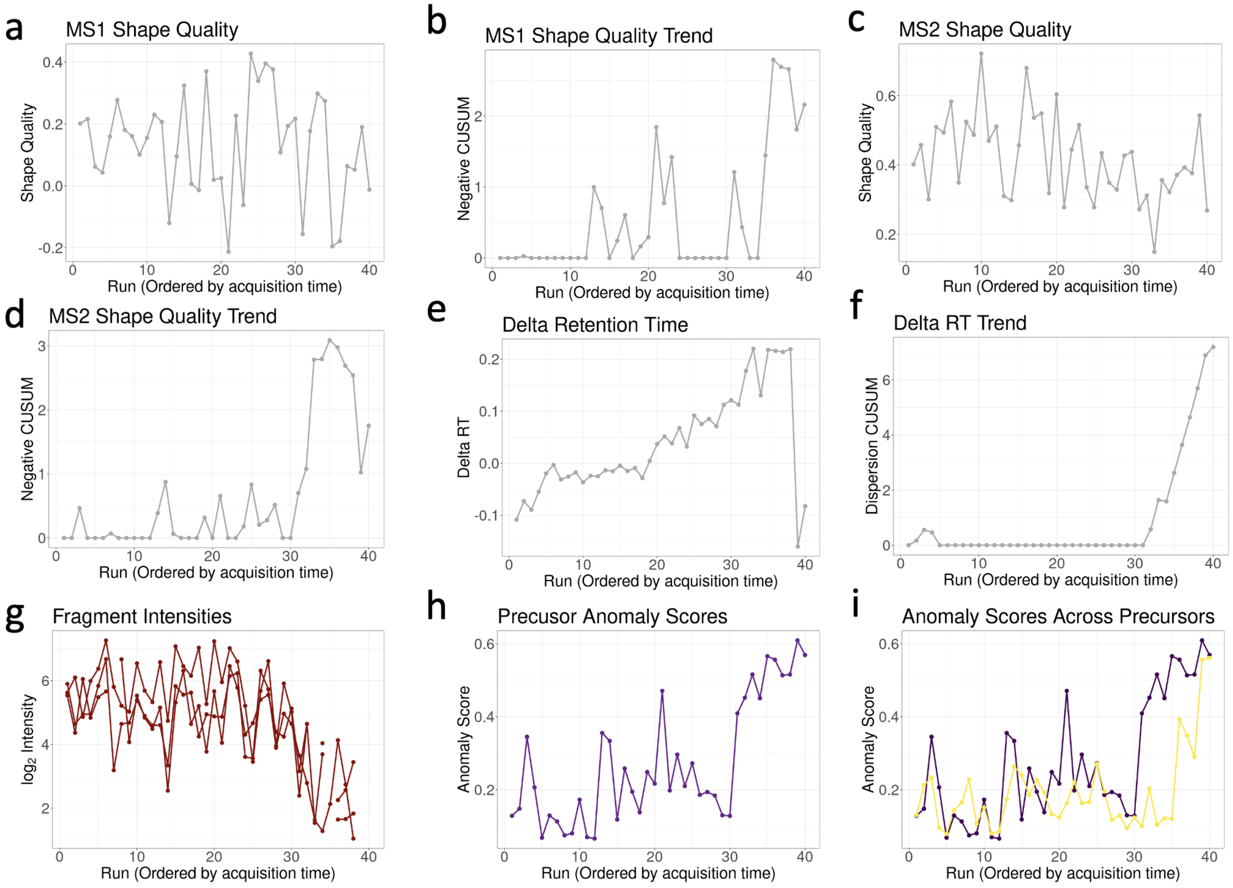

Supplementary Fig. 9: **Dataset 1: K562 benchmark. Precursor-level quality metrics for protein Q96QD8 and precursor SHYADVDPENQNFLLESNLGK.3.** *MSstats+* detected the manually introduced anomalous samples after run 30, but also identified multiple jumps in runs 3, 13, 14, and 21. (**a**) MS1 shape quality scores across all runs. Other than the known anomalous runs, clear drops occur in runs 13 and 21. (**b**) MS1 shape quality negative CUSUM. The drops seen in (a) are also identified here. (**c**) MS2 shape quality score. Other than the known anomalous runs, drops in quality are observed in runs 3, 7, 13, 14, 19, and 21. (**d**) MS2 shape quality negative CUSUM. (**e**) Delta retention time. The metric shows a clear increase over all runs until run 39, where is there is a significant drop. (**f**) Delta retention time dispersion CUSUM. (**g**) Precursor intensities, with each line representing a different fragment. (**h**) The anomaly score in each run. Drops in MS1 and MS2 quality in runs 3, 13, 14, and 21 propagate into the anomaly scores, which integrates all the quality metrics into a single value. (**i**) The anomaly score across both precursors for protein Q96QD8. Both precursors show an increase in anomaly scores after run 30.

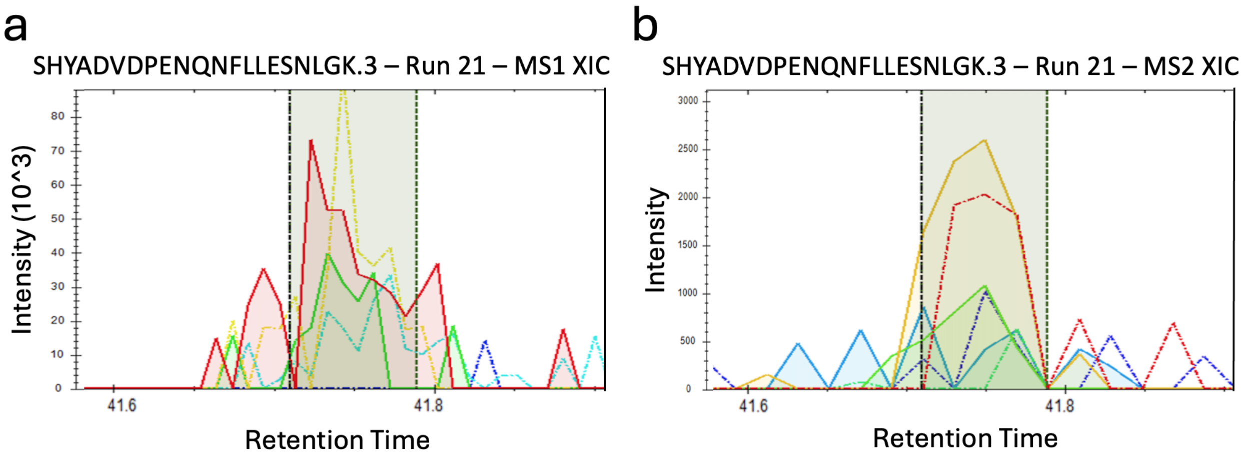

Supplementary Fig. 10: **Dataset 1: K562 benchmark. MS1 and MS2 extracted ion chromatogram (XIC) visualizations for protein Q96QD8 and precursor SHYADVDPENQNFLLESNLGK.3 in Run 21.** In **Supplementary Fig. 7**, *MSstats+* detected a singular jump in anomaly score in run 21, which can be manually validated using the XIC. The predicted retention time is indicated by the black dotted line, and the observed retention window is highlighted in green. (**a**) MS1 XIC for SHYADVDPENQNFLLESNLGK.3. Each colored line represents an isotopic peak of the precursor. The MS1 XIC peak shape was highly irregular and did not follow the expected Gaussian shape. This corresponds to the observation in **Supplementary Fig. 7**a, where the MS1 shape quality decreased significantly in run 21. (**b**) MS2 XIC for SHYADVDPENQNFLLESNLGK.3. Each colored line represents a fragment ion of the precursor. The MS2 XIC showed two fragment peaks which were generally aligned and four which appeared to be noise. While not perfect, the two aligned fragments in the MS2 XIC indicated this quantification may be reasonable. This corresponds to the observation in **Supplementary Fig. 7**c, where the MS2 shape quality was low in run 21, but not anomalously low (i.e., there were other measurements with similar MS2 shape quality).

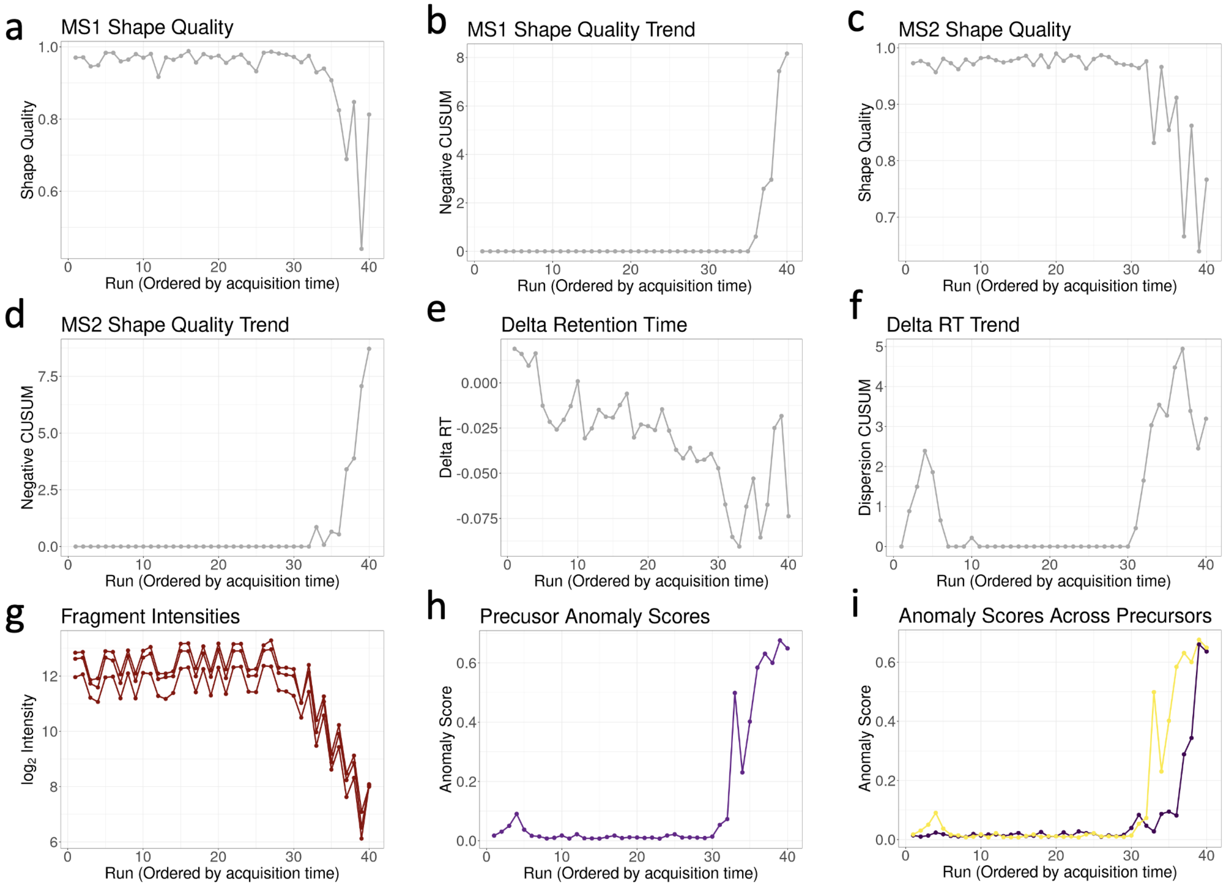

Supplementary Fig. 11: **Dataset 1: K562 benchmark. Precursor-level quality metrics for protein P55854 and precursor VAGQDGSVVQFK.2.** *MSstats+* detected the manually introduced anomalous samples starting in run 33, but was unable to identify the known anomalous runs in 31 and 32. (**a**) MS1 shape quality scores across all runs. Drops in quality began to occur after run 34. (**b**) MS1 shape quality negative CUSUM. The drops seen in (a) are also identified here. (**c**) MS2 shape quality score. Drops in quality began to occur after run 32. (**d**) MS2 shape quality negative CUSUM. (**e**) Delta retention time, which declines until it bottoms out in run 33 and then shows increased variance. (**f**) Delta retention time dispersion CUSUM. (**g**) Precursor intensities, with each line representing a different fragment. The intensities are far from the limit of detection and the drop after run 30 is clear. (**h**) The anomaly score in each run. High-quality metrics before run 30 correspond to low scores, which rise sharply with the anomalous runs. (**i**) The anomaly score across both precursors for protein Q96QD8. Both precursors show an increase in anomaly scores after run 30.

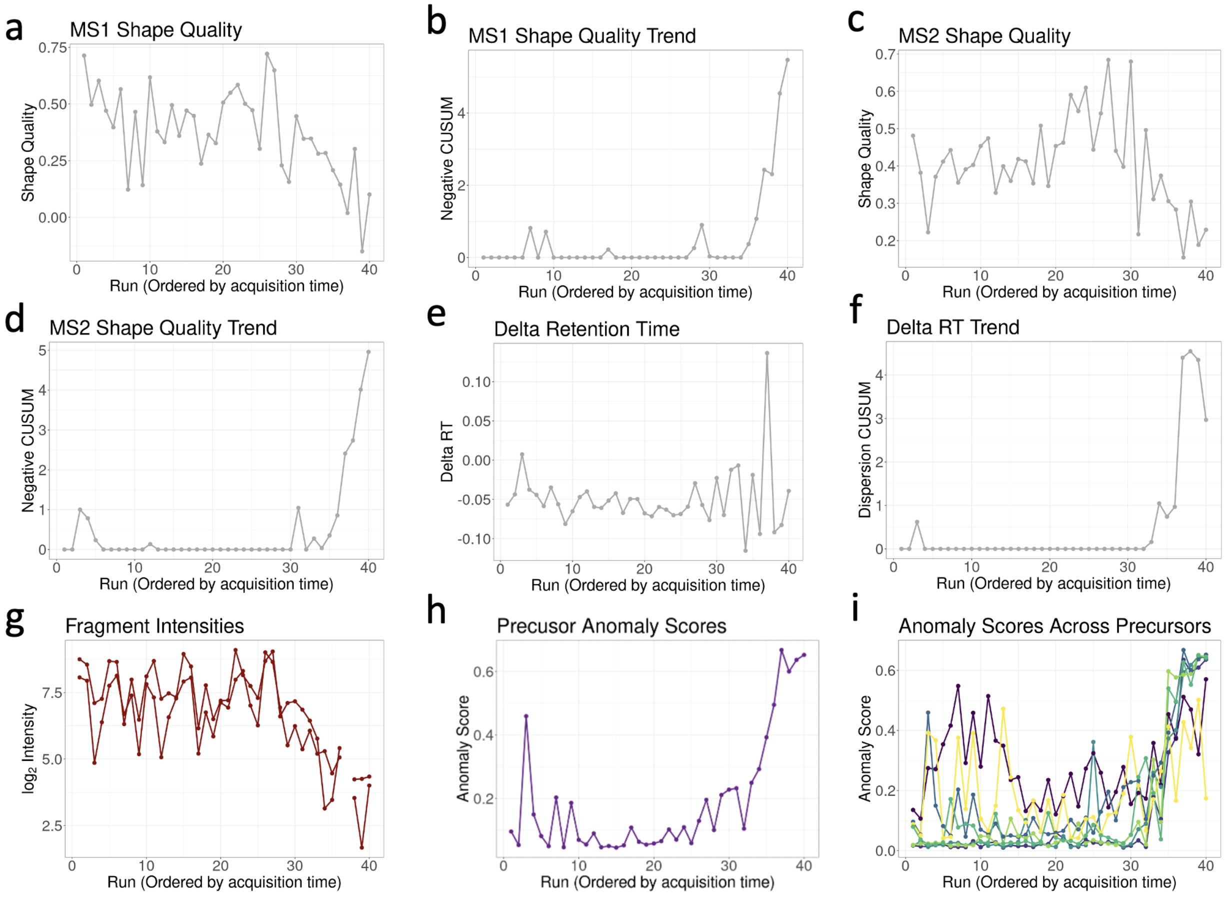

Supplementary Fig. 12: **Dataset 2: CSF benchmark. Precursor-level quality metrics for protein A0A075B7F0 and precursor LSC[Carbamidomethyl (C)]AGSGFTFSSYAM[Oxidation (M)]HWVR.3.** *MSstats+* detected the manually introduced anomalous samples starting in run 33 and identified a clear jump in run 3. (**a**) MS1 shape quality scores across all runs. Drops in quality began to occur after run 34. (**b**) MS1 shape quality negative CUSUM. (**c**) MS2 shape quality score. Drops in quality occur in the known anomalous runs, as well as in run 3. (**d**) MS2 shape quality negative CUSUM. (**e**) Delta retention time. The metric is generally stable, without any outliers, until after run 30, when the variance of the measurements increases. (**f**) Delta retention time dispersion CUSUM. (**g**) Precursor intensities, with each line representing a different fragment. (**h**) The anomaly score in each run. There is a clear jump in run 3 caused by the drop in MS2 peak shape quality. The score is then low until after run 30. (**i**) The anomaly score across all precursors for protein A0A075B7F0. Two of the precursors show an increase before run 15, and all precursors increase after run 30.

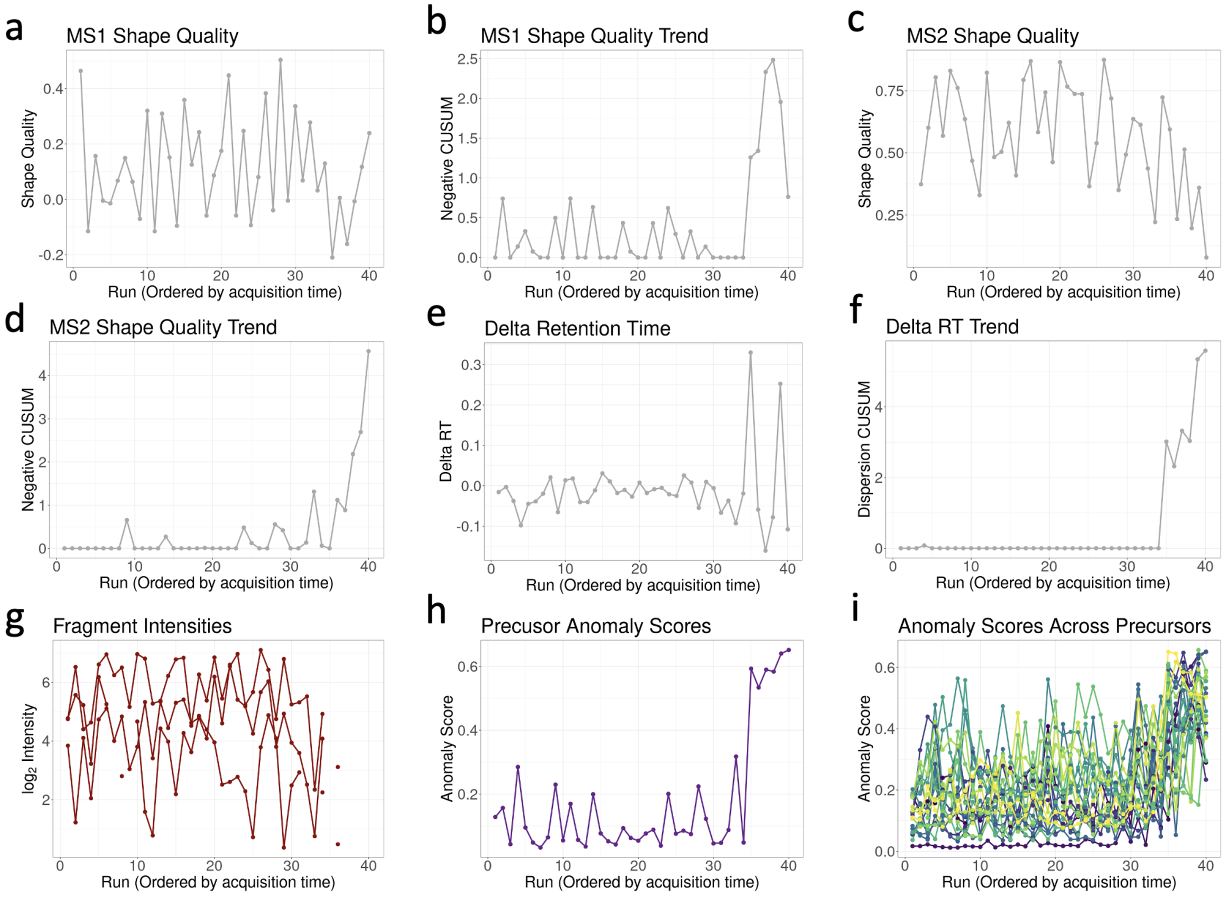

Supplementary Fig. 13: **Dataset 2: CSF benchmark. Precursor-level quality metrics for protein Q9UIW2 and precursor EVAPITR.2.** *MSstats+* detected the manually introduced anomalous samples starting in run 33 and identified several increases across the other runs. (**a**) MS1 shape quality scores across all runs. The quality was highly variable, with elevated values in some runs and significant drops in others (**b**) MS1 shape quality negative CUSUM. The variability in (a) was carried over into the CUSUM metric, with many jumps across all runs before a large increase in run 35. (**c**) MS2 shape quality score. Similar to (a) the MS2 quality was highly variable with many drops before an overall downward trend after run 30. (**d**) MS2 shape quality negative CUSUM. (**e**) Delta retention time. The metric was generally stable, without any outliers, until run 35, when the variance of the measurements notably increased. (**f**) Delta retention time dispersion CUSUM. (**g**) Precursor intensities, with each line representing a different fragment. The intensities were close to the limit of detection and exhibited increased noise and missing values. (**h**) The anomaly score in each run. The drops in quality seen in (a) and (c) are observed in small increases in anomaly score across many runs. Starting in run 33, the anomaly score increased significantly. (**i**) The anomaly score across all precursors for protein Q9UIW2. Many precursors showed jumps in anomaly score before run 30, but there was not a clear trend across all precursors until after run 30.

| Protein | Dataset | MSstats+ | MSstats | msqrob2 | limma | limpa | DEqMS |
|---|---|---|---|---|---|---|---|
| Q9Y5S9 | K562 benchmark | 0.0002 | 0.9269 | 0.8177 | 0.6134 | 0.9738 | 0.6926 |
| Q96QD8 | K562 benchmark | 0.0064 | 0.9130 | 0.7906 | 0.5247 | 0.9209 | 0.6837 |
| P55854 | K562 benchmark | 0.0001 | 0.9130 | 0.8231 | 0.6126 | 0.9196 | 0.6926 |
| A0A075B7F0 | CSF benchmark | 0.0002 | 0.6008 | 0.5476 | 0.4373 | 0.6991 | 0.5140 |
| Q9UIW2 | CSF benchmark | 0.0114 | 0.6516 | 0.5760 | 0.4107 | 0.7510 | 0.5140 |

Supplementary Table 3: Comparison of adjusted $p$-values across all statistical methods for the randomly sampled proteins across the K562 and CSF benchmark datasets in **Supplementary Fig. 7-Supplementary Fig. 13**. *MSstats+* was the only statistical method to identify any of the proteins as differentially abundant ($\alpha$=0.05). This is expected as *MSstats+* was designed to reduce the effect of the variable runs, introduced after run 30. The existing methods treated these runs as reasonable samples, increasing the standard error and resulting in elevated adjusted p-values.

#### 2.6 Coefficient of skewness distributions exhibits isolation forest model fit

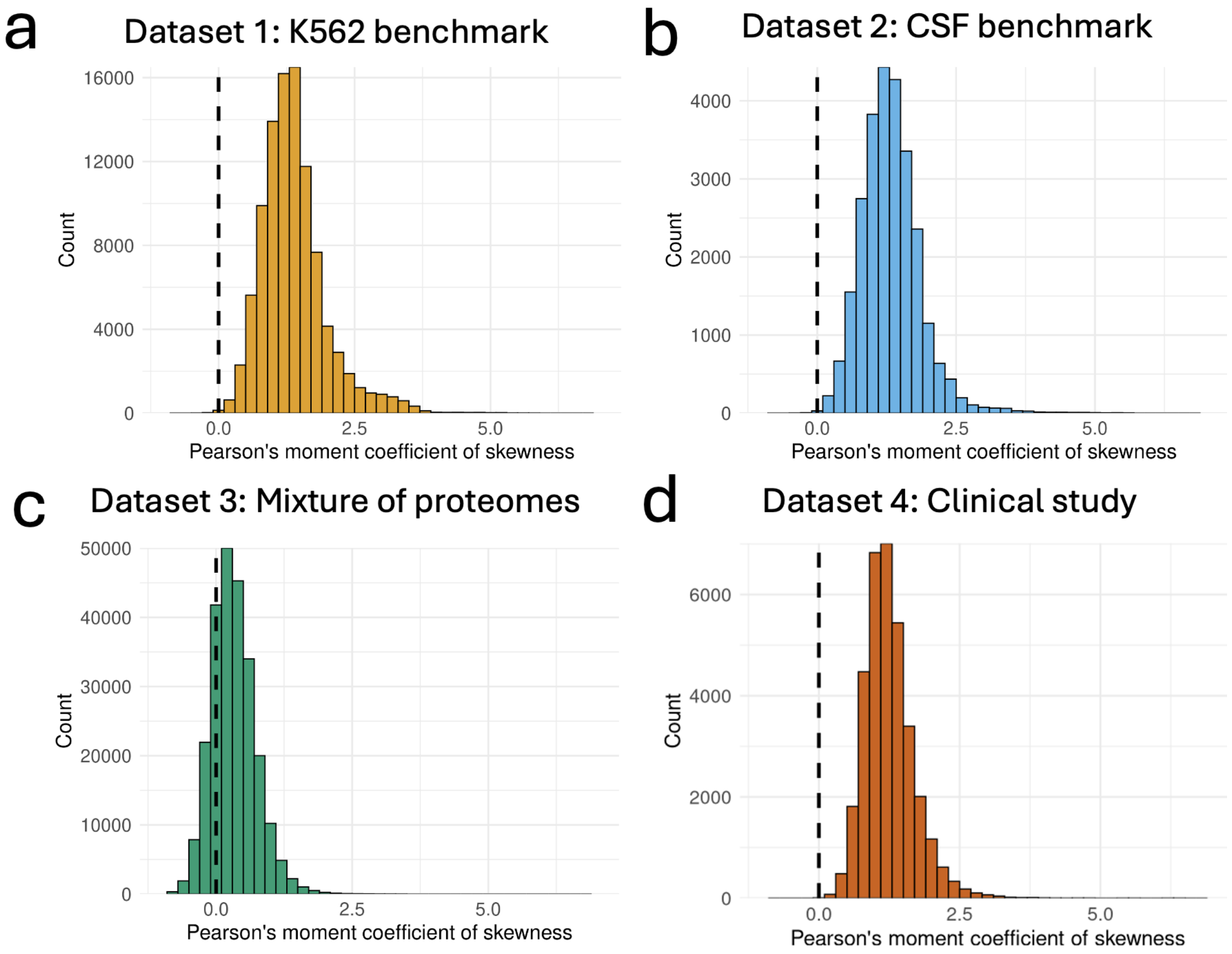

Supplementary Fig. 14: **Distribution of Pearson's moment coefficient of skewness for all precursors across the four experimental datasets.** (**a**) Dataset 1: K562 benchmark. The distribution is skewed to the right, with a median near 1 and most values above 0, indicating that the model consistently identified anomalous values across precursors. (**b**) Dataset 2: CSF benchmark. Similar (a), the distribution is shifted to the right, signaling that the model was able to detect anomalous values. (**c**) Dataset 3: Mixture of proteomes. The distribution is much closer to 0, with a median around 0.25, indicating the proposed method could not identify distinct anomalies across all precursors. This is expected given the design of the experiment. (**d**) Dataset 4: Clinical cohort. The distribution is shifted right with a median near 1, highlighting the proposed methods effectiveness in real world clinical studies.

#### 2.7 Anomaly scores for known anomalous and non-anomalous runs shows clearly different distributions

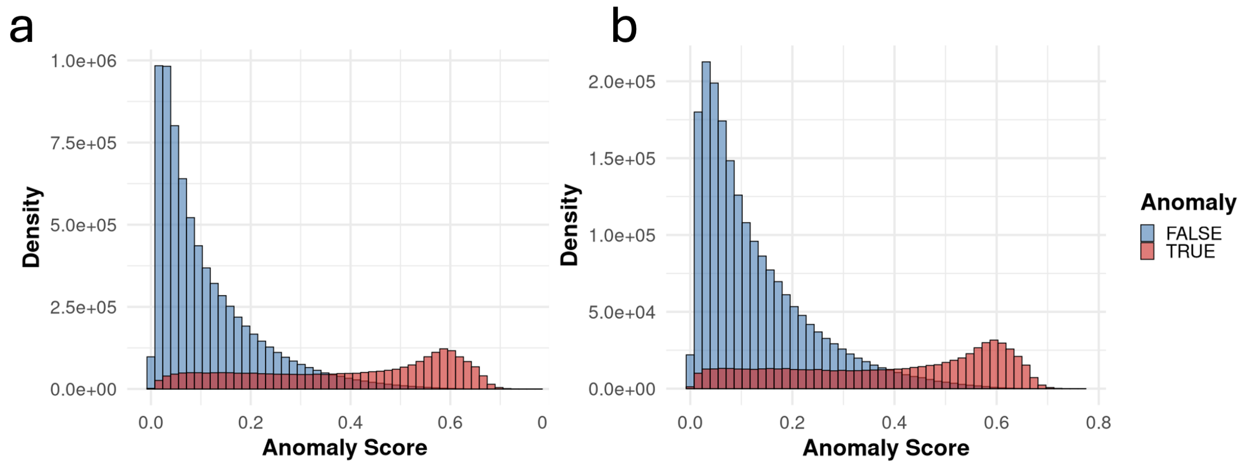

Supplementary Fig. 15: ***MSstats+* is able to differentiate between known anomalous and non-anomalous runs in two benchmark experiments.** **(a)** Dataset 1: K562 benchmark. Distribution of anomaly scores across all precursors subset by anomalous (red) and non-anomalous (blue) runs. There was a clear separation between the manually introduced anomalous runs and non-anomalous runs. Non-anomalous runs exhibit a right shifted distribution with most scores close to zero and a long tail towards higher scores. In contrast, anomalous runs were left shifted with a peak around 0.6 and a broader spread towards smaller scores. Non-anomalous runs may still contain low-quality measurements due to a variety of factors (e.g., poor ionization efficiency or low signal). Likewise, anomalous runs, especially those which are towards the start of the manually introduced instrumental drift, may exhibit patterns that resemble higher quality measurements leading to lower scores. **(b)** Same as (a) for Dataset 2: CSF benchmark. The observations in (a) hold for the CSF benchmark, with slightly higher anomaly scores for the non-anomalous runs likely driven by the increased complexity in the samples.

#### 2.8 Measurement anomaly scores are not exactly correlated with intensity values

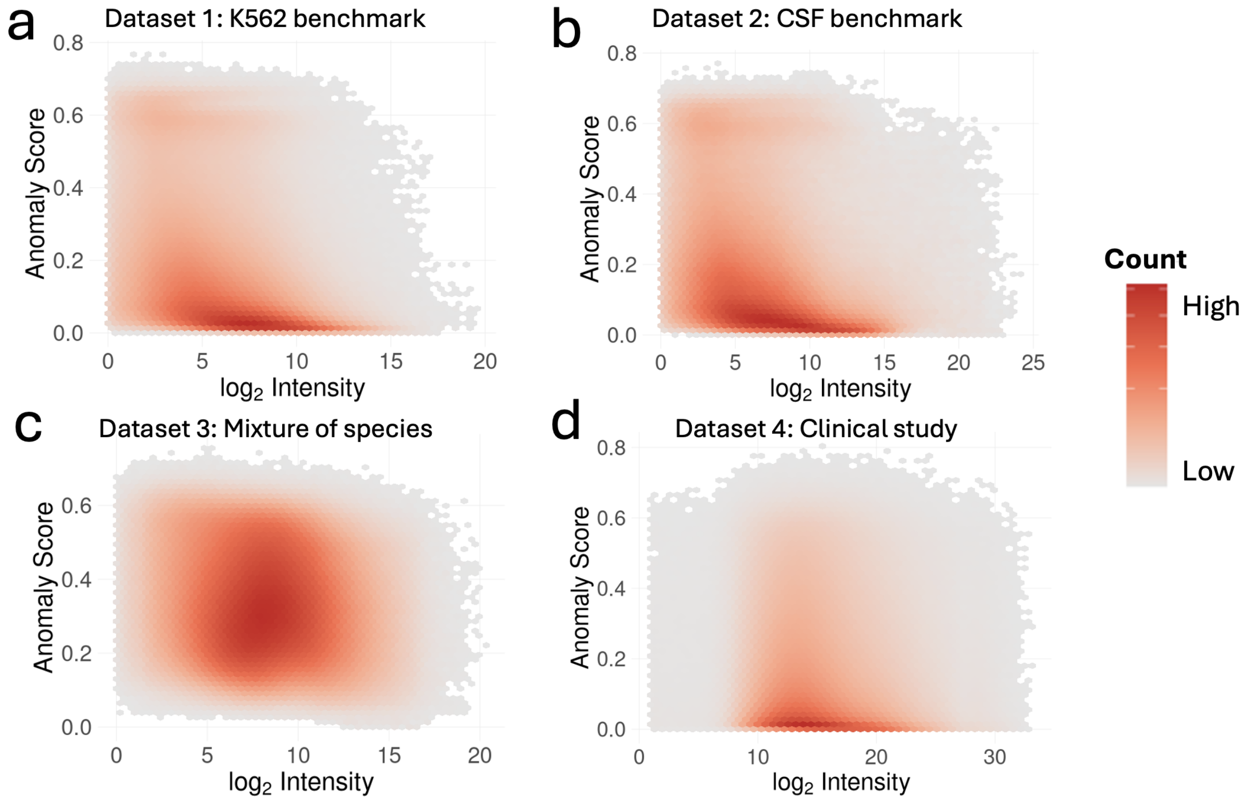

Supplementary Fig. 16: **$\mathbf{Log_2}$ intensities and anomaly scores do not show high correlation across the four experimental datasets.** (**a**) Dataset 1: K562 benchmark. Most anomaly scores were low across the full intensity range and there was a slight negative trend with lower intensities tending to have higher anomaly scores. However, there were still many low intensities with low anomaly scores and higher intensities with high anomaly scores. (**b**) Dataset 2: CSF benchmark. The correlations exhibited similar patterns as in (a), with more anomaly scores in the mid-range ( 0.2-0.4) likely driven by the additional complexity in the biological material used. (**c**) Mixture of proteomes dataset. The correlation did not exhibit the same patterns seen in (a) and (b). Instead of the loose negative exponential shape, the distribution was mostly uncorrelated. This lack of correlation is expected given the distribution of skewed coefficients seen in **Supplementary Fig. 14**. (**d**) Clinical study dataset. The correlations followed a similar trend to (a) and (b) but the overall intensity was shifted away from zero. There were many fragments which measured low intensities and showed low anomaly scores and vice versa.

##### 2.9 Additional peak quality metrics for AAAGGPGGAALGEAPPGR.2 - Q6UW01 highlight samples with lower quality

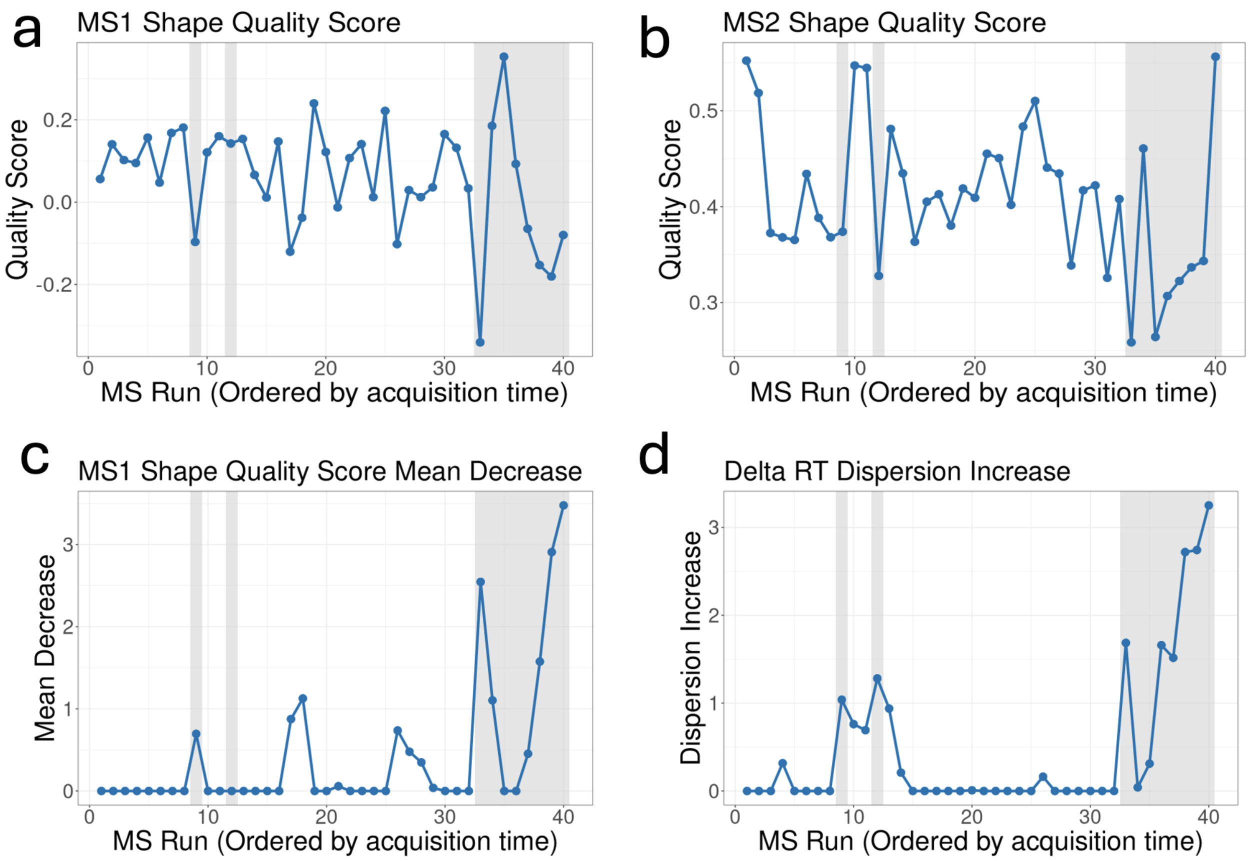

Supplementary Fig. 17: **Dataset 2: CSF benchmark. Precursor-level quality metrics for protein Q6UW01 and precursor AAAGGPGGAALGEAPPGR.2 show a drop in quality in runs 9, 12, and 33-40. Elevated anomaly scores ($> 0.35$) are highlighted in gray.** (**a**) The MS1 shape quality scores across all runs. The MS1 shape quality score sees a substantial drop in run 9 compared to the average in previous runs, and increased variability across runs 33-40. (**b**) Same as (a) for MS2 shape quality score. The MS2 shape quality score is lower in runs 9 and 12, and shows a particularly stark drop between runs 11 and 12. Again in runs 33-40 there is increased variability. (**c**) Same as (a) for the engineered MS1 shape quality negative CUSUM metric. This metric captures downward trends in shape quality score at the MS1 level. The drop in run 9 is captured by this metric, along with the increased variability in runs 33-40. (**d**) Same as (a) for the engineered delta retention time variance CUSUM metric and captures variation in the delta retention time. This metric shows an area of increased scores from runs 9 to 13, indicating an area of samples where the delta retention time was unstable. Runs 33-40 again show increased variability.

#### 2.10 Anomaly scores act as a natural proxy for confidence in missing value imputation

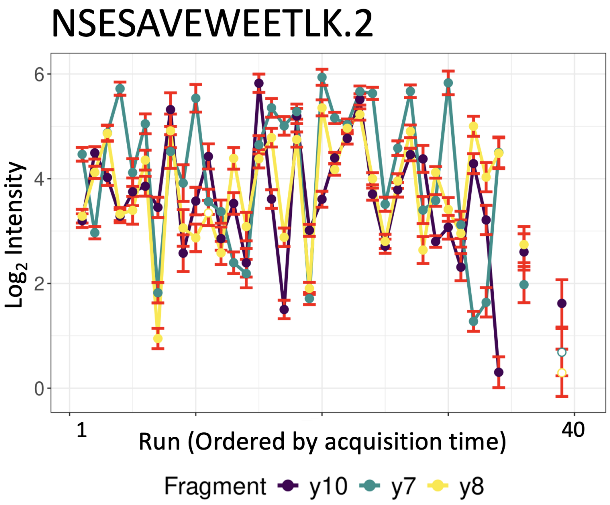

Supplementary Fig. 18: **Dataset 1: K562 benchmark. Profile plot for precursor NSESAVEWEETLK.2 in protein Q9NWH2 highlighting where imputation is not performed.** Profile plot with temporally ordered runs are on the x-axis and $\log_2$ intensity is on the y-axis. Each colored line represents a different fragment ion for the precursor. Solid points indicate observed values, and hollow (white) points denote imputed values. Red error bars represent the anomaly scores in each run. In run 39 the proposed method imputed the y7 and y8 fragments. in contrast, in runs 35, 37, 38, and 40 imputation was not performed for any fragment. The proposed approach will only perform imputation if there is at least one observed measurement for the protein in the run (across all precursors). There were no observed measurements for runs 35, 37, 38, or 40 and imputation was not performed due to lack of information.

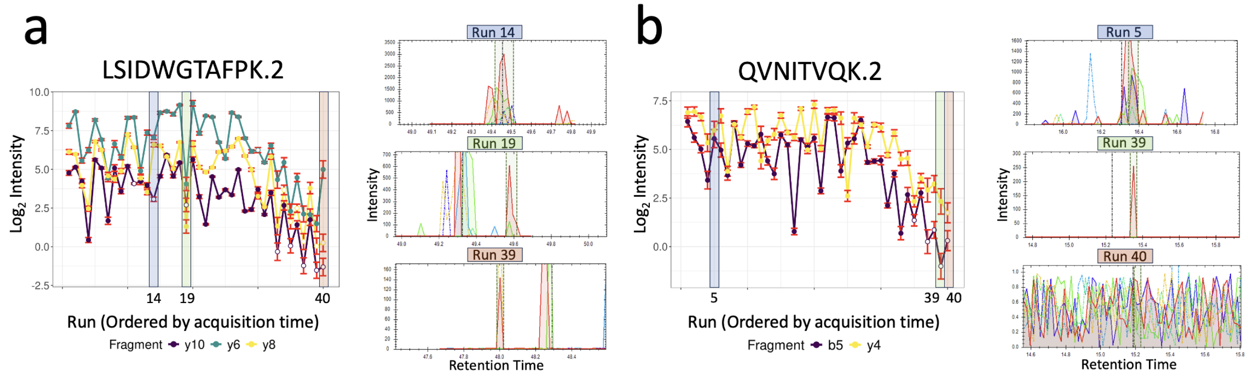

Supplementary Fig. 19: **Dataset 1: K562 benchmark: the underlying peak quality metrics provide a quantitative measure for confidence in the effectiveness of missing value imputation.** For both precursors, temporally ordered runs are on the x-axis and $\log_2$ intensity is on the y-axis. Each colored line represents a different fragment ion for the precursor. Solid points indicate observed values, and hollow (white) points denote imputed values. Red error bars represent the anomaly scores in each run. Plots on the right show the corresponding spectral peaks for selected runs. **(a)** Precursor LSIDWGTAFPK.2 for protein Q9BZG8, runs 14, 19, and 40 are highlighted. In run 14, the y10 ion was missing, however the shape of the observed peaks was reasonable resulting in a low anomaly score and high confidence in the imputation. In contrast, in run 19 y10 was also missing, however the observed peak shape exhibited lower quality, resulting in a high anomaly score and low confidence in the imputation. Finally, in run 40 both y10 and y8 were imputed. The observed peak quality was low and the resulting imputations had high anomaly scores and low confidence. **(b)** Precursor QVNITVQK.2 for protein Q9P035, runs 5, 39, and 40 are highlighted. In run 5, both fragments were observed and no imputation was required. In run 39 the b5 ion was imputed and the y4 ion was observed (shown as a single peak). The peak quality for y4 was low, resulting in low imputation confidence. Finally, in run 40 both the b6 and y4 ions were imputed (only noise was observed). Imputation could be performed as other precursors were measured in run 40 for protein Q9P035, however due to the low peak quality, the anomaly score was high and the imputation confidence was low.

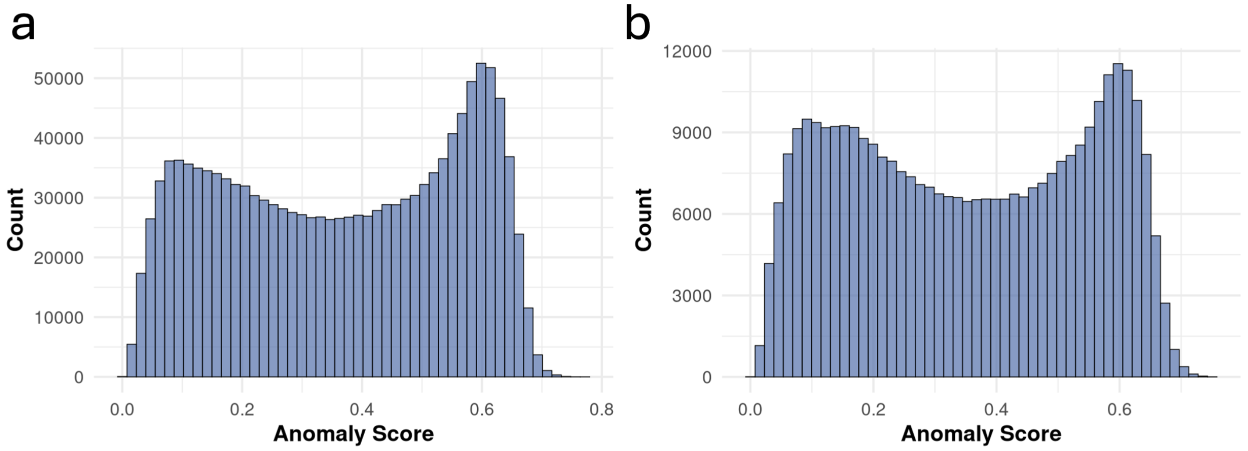

Supplementary Fig. 20: **Imputed measurements have an anomaly score distribution shifted towards higher scores. (a)** Dataset 1: K562 benchmark. **(c)** Dataset 2: CSF benchmark. Across both datasets, imputed values exhibit a bimodal distribution, with a low peak near 0 and a higher peak around 0.6. This is in stark contrast to observed anomaly scores whose distribution was right skewed (**Supplementary Fig. 15**). These distributions highlight that not all imputations are treated uniformly in protein-level summarization. Instead their weights are determined by the anomaly score, reflecting the underlying peak quality.

#### 2.11 *MSstats+* outperforms existing statistical methods in terms of FDR on a controlled mixture of proteomes

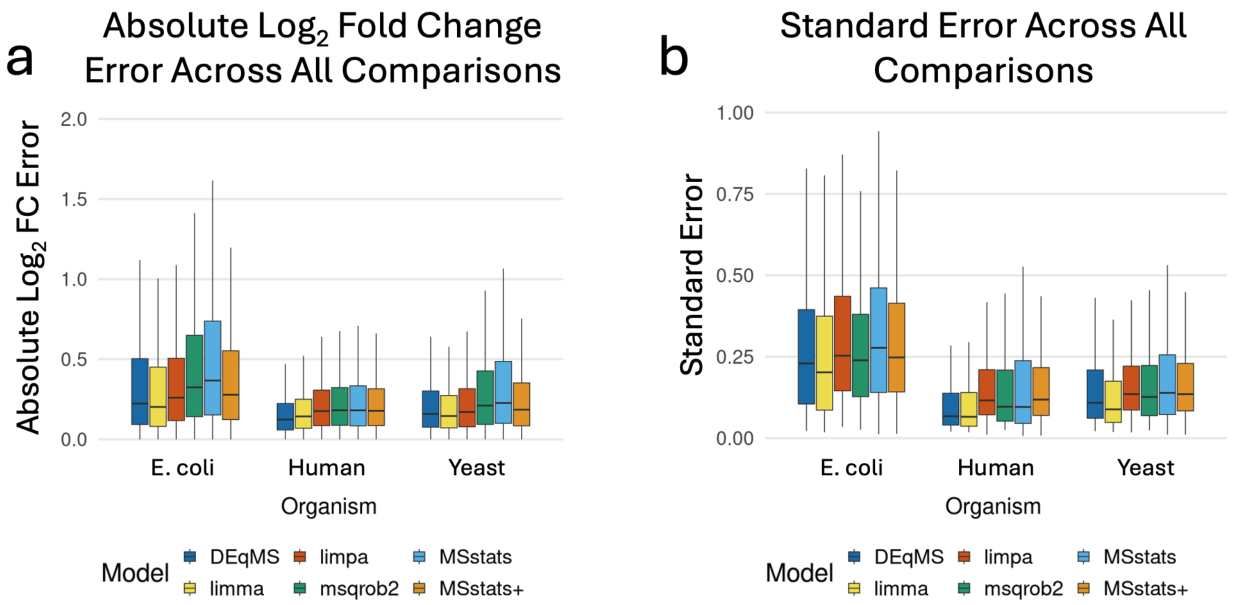

Supplementary Fig. 21: **Dataset 3: Mixture of proteomes: *MSstats+* estimates a comparable fold change while avoiding overconfident standard errors.** (**a**) The absolute difference between the estimated fold change and the ground truth fold change across all proteins and comparisons for each model. *MSstats+* produced fold change errors that were broadly consistent with other statistical models. (**b**) The standard error across all proteins and comparisons for each model. *MSstats+* balanced between having low standard errors while avoiding unrealistically small standard errors, reducing the risk of false positives.

#### 2.12 MSstats+ controls for poor quantitative measurements and reduces variation in a real world clinical study compared to MSstats

| Model | $Log_2$ FC | SE | DF | Adj. p-value |
|---|---|---|---|---|
| MSstats+ | 0.685 | 0.179 | 144 | 0.017 |
| MSstats | 0.652 | 0.222 | 144 | 0.254 |

Supplementary Table 5: **Comparison of differential analysis results for P11597 (CETP) in Dataset 4: Clinical study between *MSstats+* and *MSstats*.** Both methods estimated a similar $\log_2$ fold change, but *MSstats+* downweighted poor-quality measurements, resulting in a reduced standard error and smaller adjusted p-value.

### 3 Experimental data

| Dataset | Conditions | Replicates | Proteins | Avg. Peptides / Protein | Avg. Fragments / Peptide |
|---|---|---|---|---|---|
| 1: K562 benchmark | 2 | 20 | 6382 | 14.26 | 2.67 |
| 2: CSF benchmark | 2 | 20 | 2529 | 9.37 | 2.74 |
| 3: Mixture of proteomes | 6 | 3 | 14257 | 13.89 | 3.20 |
| 4: Clinical cohort | 2 | 123 | 3184 | 10.85 | 2.94 |

Supplementary Table 6: **Experimental design and measurement overview for the experimental datasets included in this manuscript.** Each dataset represents a distinct experimental design. The number of replicates represents the average number of replicates per condition. The number of proteins reflects filtered protein groups after quality control and preprocessing. Peptide and fragment counts are averaged across all proteins.

#### 3.1 Experimental design of Dataset 1: K562 benchmark & Dataset 2: CSF benchmark

In both Dataset 1 and Dataset 2, all proteins were differentially abundant by default (due to the experimental design). To mimic a real world scenario, we randomly selected 90% of the proteins in each experiment and swapped the condition labels, such that they would no longer be differentially abundant (expected $\log_2$ fold change of 0). The exact condition swaps (i.e., which runs were swapped and which stayed the same) are listed in the tables below.

##### 3.1.1 Dataset 1: K562 benchmark

| Run Name | Ratio | Original Condition | Swapped Condition |
|---|---|---|---|
| 20250102_Tulum_NeatK562_Seq1 | Neat | Condition1 | Condition1 |
| 20250102_Tulum_NeatK562_Seq2 | Neat | Condition1 | Condition2 |
| 20250102_Tulum_1to2K562_Seq3 | 1:2 | Condition2 | Condition2 |
| 20250102_Tulum_1to2K563_Seq4 | 1:2 | Condition2 | Condition1 |
| 20250102_Tulum_NeatK562_Seq5 | Neat | Condition1 | Condition1 |
| 20250102_Tulum_NeatK562_Seq6 | Neat | Condition1 | Condition2 |
| 20250102_Tulum_1to2K562_Seq7 | 1:2 | Condition2 | Condition2 |
| 20250102_Tulum_NeatK562_Seq8 | Neat | Condition1 | Condition1 |
| 20250102_Tulum_1to2K562_Seq9 | 1:2 | Condition2 | Condition1 |
| 20250102_Tulum_NeatK562_Seq10 | Neat | Condition1 | Condition2 |
| 20250102_Tulum_NeatK562_Seq11 | Neat | Condition1 | Condition1 |
| 20250102_Tulum_1to2K562_Seq12 | 1:2 | Condition2 | Condition2 |
| 20250102_Tulum_1to2K562_Seq13 | 1:2 | Condition2 | Condition1 |
| 20250102_Tulum_1to2K562_Seq14 | 1:2 | Condition2 | Condition2 |
| 20250102_Tulum_NeatK562_Seq15 | Neat | Condition1 | Condition2 |
| 20250102_Tulum_NeatK562_Seq16 | Neat | Condition1 | Condition1 |
| 20250102_Tulum_1to2K562_Seq17 | 1:2 | Condition2 | Condition1 |
| 20250102_Tulum_NeatK562_Seq18 | Neat | Condition1 | Condition2 |
| 20250102_Tulum_1to2K562_Seq19 | 1:2 | Condition2 | Condition2 |
| 20250102_Tulum_NeatK562_Seq20 | Neat | Condition1 | Condition1 |
| 20250102_Tulum_1to2K562_Seq21 | 1:2 | Condition2 | Condition1 |
| 20250102_Tulum_NeatK562_Seq22 | Neat | Condition1 | Condition2 |
| 20250102_Tulum_NeatK562_Seq23 | Neat | Condition1 | Condition1 |
| 20250102_Tulum_1to2K562_Seq24 | 1:2 | Condition2 | Condition2 |
| 20250102_Tulum_1to2K562_Seq25 | 1:2 | Condition2 | Condition1 |
| 20250102_Tulum_NeatK562_Seq26 | Neat | Condition1 | Condition2 |
| 20250102_Tulum_NeatK562_Seq27 | Neat | Condition1 | Condition1 |
| 20250102_Tulum_1to2K562_Seq28 | 1:2 | Condition2 | Condition2 |
| 20250102_Tulum_1to2K562_Seq29 | 1:2 | Condition2 | Condition1 |
| 20250102_Tulum_1to2K562_Seq30 | 1:2 | Condition2 | Condition2 |
| 20250102_Tulum_1to4K562_Seq31 | 1:4 | Condition2 | Condition1 |
| 20250102_Tulum_1to2K562_Seq32 | 1:2 | Condition1 | Condition2 |
| 20250102_Tulum_1to8K562_Seq33 | 1:8 | Condition2 | Condition2 |
| 20250102_Tulum_1to4K562_Seq34 | 1:4 | Condition1 | Condition1 |
| 20250102_Tulum_1to16K562_Seq35 | 1:16 | Condition2 | Condition1 |
| 20250102_Tulum_1to8K562_Seq36 | 1:8 | Condition1 | Condition2 |
| 20250102_Tulum_1to32K562_Seq37 | 1:32 | Condition2 | Condition2 |
| 20250102_Tulum_1to16K562_Seq38 | 1:16 | Condition1 | Condition1 |
| 20250102_Tulum_1to64K562_Seq39 | 1:64 | Condition2 | Condition1 |
| 20250102_Tulum_1to32K562_Seq40 | 1:32 | Condition1 | Condition2 |

Supplementary Table 7: Condition swapping key for random 90% sampled proteins in the K562 benchmark experiment. The swapping was performed such that the expected $\log_2$ fold change would be exactly 0.

##### 3.1.2 Dataset 2: CSF benchmark

| Run Name | Ratio | Original Condition | Swapped Condition |
|---|---|---|---|
| 20250123_Tulum_NeatCSF-DD_Seq1 | Neat | Condition1 | Condition1 |
| 20250123_Tulum_NeatCSF-DD_Seq2 | Neat | Condition1 | Condition2 |
| 20250123_Tulum_1to2CSF-DD_Seq3 | 1:2 | Condition2 | Condition2 |
| 20250123_Tulum_1to2CSF-DD_Seq4 | 1:2 | Condition2 | Condition1 |
| 20250123_Tulum_NeatCSF-DD_Seq5 | Neat | Condition1 | Condition1 |
| 20250123_Tulum_NeatCSF-DD_Seq6 | Neat | Condition1 | Condition2 |
| 20250123_Tulum_1to2CSF-DD_Seq7 | 1:2 | Condition2 | Condition2 |
| 20250123_Tulum_NeatCSF-DD_Seq8 | Neat | Condition1 | Condition1 |
| 20250123_Tulum_1to2CSF-DD_Seq9 | 1:2 | Condition2 | Condition1 |
| 20250123_Tulum_NeatCSF-DD_Seq10 | Neat | Condition1 | Condition2 |
| 20250123_Tulum_NeatCSF-DD_Seq11 | Neat | Condition1 | Condition1 |
| 20250123_Tulum_1to2CSF-DD_Seq12 | 1:2 | Condition2 | Condition2 |
| 20250123_Tulum_1to2CSF-DD_Seq13 | 1:2 | Condition2 | Condition1 |
| 20250123_Tulum_1to2CSF-DD_Seq14 | 1:2 | Condition2 | Condition2 |
| 20250123_Tulum_NeatCSF-DD_Seq15 | Neat | Condition1 | Condition2 |
| 20250123_Tulum_NeatCSF-DD_Seq16 | Neat | Condition1 | Condition1 |
| 20250123_Tulum_1to2CSF-DD_Seq17 | 1:2 | Condition2 | Condition1 |
| 20250123_Tulum_NeatCSF-DD_Seq18 | Neat | Condition1 | Condition2 |
| 20250123_Tulum_1to2CSF-DD_Seq19 | 1:2 | Condition2 | Condition2 |
| 20250123_Tulum_NeatCSF-DD_Seq20 | Neat | Condition1 | Condition1 |
| 20250123_Tulum_1to2CSF-DD_Seq21 | 1:2 | Condition2 | Condition1 |
| 20250123_Tulum_NeatCSF-DD_Seq22 | Neat | Condition1 | Condition2 |
| 20250123_Tulum_NeatCSF-DD_Seq23 | Neat | Condition1 | Condition1 |
| 20250123_Tulum_1to2CSF-DD_Seq24 | 1:2 | Condition2 | Condition2 |
| 20250123_Tulum_1to2CSF-DD_Seq25 | 1:2 | Condition2 | Condition1 |
| 20250123_Tulum_NeatCSF-DD_Seq26 | Neat | Condition1 | Condition2 |
| 20250123_Tulum_NeatCSF-DD_Seq27 | Neat | Condition1 | Condition1 |
| 20250123_Tulum_1to2CSF-DD_Seq28 | 1:2 | Condition2 | Condition2 |
| 20250123_Tulum_1to2CSF-DD_Seq29 | 1:2 | Condition2 | Condition1 |
| 20250123_Tulum_1to2CSF-DD_Seq30 | 1:2 | Condition2 | Condition2 |
| 20250123_Tulum_1to4CSF-DD_Seq31 | 1:4 | Condition2 | Condition1 |
| 20250123_Tulum_1to2CSF-DD_Seq32 | 1:2 | Condition1 | Condition2 |
| 20250123_Tulum_1to8CSF-DD_Seq33 | 1:8 | Condition2 | Condition2 |
| 20250123_Tulum_1to4CSF-DD_Seq34 | 1:4 | Condition1 | Condition1 |
| 20250123_Tulum_1to16CSF-DD_Seq35 | 1:16 | Condition2 | Condition1 |
| 20250123_Tulum_1to8CSF-DD_Seq36 | 1:8 | Condition1 | Condition2 |
| 20250123_Tulum_1to32CSF-DD_Seq37 | 1:32 | Condition2 | Condition2 |
| 20250123_Tulum_1to16CSF-DD_Seq38 | 1:16 | Condition1 | Condition1 |
| 20250123_Tulum_1to64CSF-DD_Seq39 | 1:64 | Condition2 | Condition1 |
| 20250123_Tulum_1to32CSF-DD_Seq40 | 1:32 | Condition1 | Condition2 |

Supplementary Table 8: Condition swapping key for random 90% sampled proteins in the CSF benchmark experiment. The swapping was performed such that the expected $\log_2$ fold change would be exactly 0.

##### 3.2 Separate replicates in Dataset 4: Clinical study into two conditions by CDR-SB score

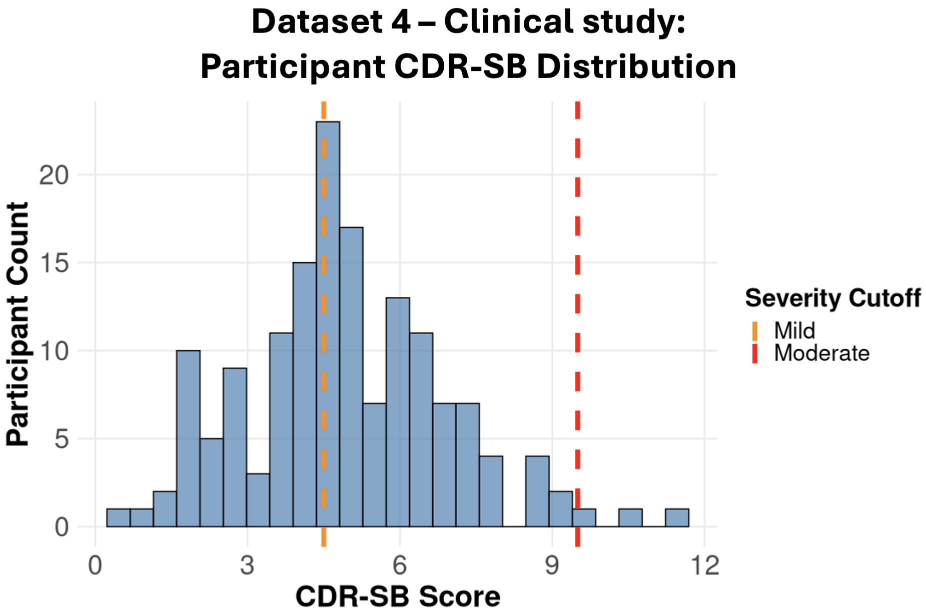

Supplementary Fig. 22: **Dataset 4: Clinical study. Participant CDR-SB score of 4.5 was chosen to separate the study into two conditions: low and high CDR-SB.** The dotted lines represent mild (orange) and moderate (red) severity cutoffs in Alzheimer's disease progression as defined by O'Bryant et al.[2]. The mild cutoff (orange) at 4.5 was chosen to ensure a similar number of participants per group.

| Label | MSstats+ | | | MSstats | | | msqrob2 | | | limma | | | limpa | | | DEqMS | | |
|---|---|---|---|---|---|---|---|---|---|---|---|---|---|---|---|---|---|---|
| | $\log_2$FC | SE | $p_{adj}$ | $\log_2$FC | SE | $p_{adj}$ | $\log_2$FC | SE | $p_{adj}$ | $\log_2$FC | SE | $p_{adj}$ | $\log_2$FC | SE | $p_{adj}$ | $\log_2$FC | SE | $p_{adj}$ |
| E10H50Y40 vs E20H50Y30 | −0.114 | 0.179 | 0.705 | −0.050 | 0.159 | 0.834 | −0.046 | 0.157 | 0.848 | −0.096 | 0.102 | 0.486 | −0.077 | 0.133 | 0.710 | −0.147 | 0.103 | 0.317 |
| E10H50Y40 vs E30H50Y20 | 0.253 | 0.175 | 0.239 | 0.322 | 0.159 | 0.091 | 0.291 | 0.157 | 0.116 | 0.304 | 0.102 | 0.016 | 0.301 | 0.135 | 0.047 | 0.272 | 0.103 | 0.032 |
| E10H50Y40 vs E40H50Y10 | −0.054 | 0.173 | 0.851 | −0.112 | 0.159 | 0.633 | −0.088 | 0.157 | 0.705 | −0.022 | 0.102 | 0.897 | −0.001 | 0.131 | 0.996 | 0.026 | 0.103 | 0.891 |
| E10H50Y40 vs E45H50Y5 | −0.036 | 0.176 | 0.894 | −0.198 | 0.159 | 0.353 | −0.137 | 0.157 | 0.506 | −0.014 | 0.102 | 0.930 | −0.045 | 0.130 | 0.808 | −0.019 | 0.103 | 0.914 |
| E10H50Y40 vs E5H50Y45 | 0.347 | 0.180 | 0.184 | 0.457 | 0.159 | 0.042 | 0.332 | 0.159 | 0.122 | 0.271 | 0.102 | 0.046 | 0.361 | 0.138 | 0.044 | 0.245 | 0.103 | 0.085 |
| E20H50Y30 vs E30H50Y20 | 0.367 | 0.169 | 0.079 | 0.372 | 0.159 | 0.055 | 0.336 | 0.153 | 0.065 | 0.400 | 0.102 | 0.003 | 0.378 | 0.136 | 0.013 | 0.418 | 0.103 | 0.003 |
| E20H50Y30 vs E40H50Y10 | 0.060 | 0.166 | 0.812 | −0.062 | 0.159 | 0.801 | −0.042 | 0.153 | 0.847 | 0.074 | 0.102 | 0.587 | 0.076 | 0.132 | 0.678 | 0.172 | 0.103 | 0.202 |
| E20H50Y30 vs E45H50Y5 | 0.078 | 0.169 | 0.733 | −0.148 | 0.159 | 0.460 | −0.091 | 0.153 | 0.633 | 0.082 | 0.102 | 0.506 | 0.032 | 0.131 | 0.853 | 0.127 | 0.103 | 0.317 |
| E20H50Y30 vs E5H50Y45 | 0.460 | 0.173 | 0.039 | 0.507 | 0.159 | 0.014 | 0.378 | 0.155 | 0.045 | 0.367 | 0.102 | 0.006 | 0.438 | 0.138 | 0.006 | 0.391 | 0.103 | 0.005 |
| E30H50Y20 vs E40H50Y10 | −0.307 | 0.162 | 0.134 | −0.433 | 0.159 | 0.030 | −0.379 | 0.153 | 0.044 | −0.326 | 0.102 | 0.012 | −0.302 | 0.134 | 0.049 | −0.246 | 0.103 | 0.053 |
| E30H50Y20 vs E45H50Y5 | −0.289 | 0.165 | 0.172 | −0.520 | 0.159 | 0.012 | −0.427 | 0.153 | 0.026 | −0.318 | 0.102 | 0.014 | −0.346 | 0.133 | 0.023 | −0.291 | 0.103 | 0.025 |
| E30H50Y20 vs E5H50Y45 | 0.094 | 0.169 | 0.696 | 0.135 | 0.159 | 0.505 | 0.042 | 0.155 | 0.845 | −0.033 | 0.102 | 0.802 | 0.060 | 0.140 | 0.755 | −0.027 | 0.103 | 0.845 |
| E40H50Y10 vs E45H50Y5 | 0.018 | 0.162 | 0.957 | −0.087 | 0.159 | 0.757 | −0.049 | 0.153 | 0.853 | 0.008 | 0.102 | 0.971 | −0.044 | 0.129 | 0.846 | −0.045 | 0.103 | 0.836 |
| E40H50Y10 vs E5H50Y45 | 0.401 | 0.166 | 0.059 | 0.568 | 0.159 | 0.007 | 0.420 | 0.155 | 0.030 | 0.293 | 0.102 | 0.021 | 0.362 | 0.136 | 0.021 | 0.219 | 0.103 | 0.083 |
| E45H50Y5 vs E5H50Y45 | 0.383 | 0.170 | 0.082 | 0.655 | 0.159 | 0.003 | 0.469 | 0.155 | 0.020 | 0.285 | 0.102 | 0.027 | 0.406 | 0.135 | 0.010 | 0.264 | 0.103 | 0.045 |

Supplementary Table 4: **Dataset 3: Mixture of proteomes. Model estimates for $\log_2$ fold-change, standard error, and adjusted p-value across different comparisons and statistical methods for protein O15091.** O15091 is a human protein and is expected to have a log2 fold change of zero in all comparisons with an adjusted p-value above 0.05. *MSstats+* identified only one comparison as a false positive, with adjusted p-values above 0.05. In comparison, all other methods identified at least five false positives.
